# Biosynthesis of iridoid sex pheromones in aphids

**DOI:** 10.1101/2022.09.13.507830

**Authors:** Tobias G. Köllner, Anja David, Katrin Luck, Franziska Beran, Grit Kunert, Jing-Jiang Zhou, Lorenzo Caputi, Sarah E. O’Connor

## Abstract

Iridoid monoterpenes, widely distributed in plants and insects, have many ecological functions. While the biosynthesis of iridoids has been extensively studied in plants, little is known about how insects synthesize these natural products. Here, we elucidated the biosynthesis of the iridoids *cis*-*trans-*nepetalactol and *cis*-*trans-*nepetalactone in the pea aphid *Acyrthosiphon pisum* [Harris], where they act as sex pheromones. The exclusive production of iridoids in hind legs of sexual female aphids allowed us to identify iridoid genes by searching for genes specifically expressed in this tissue. Biochemical characterization of candidate enzymes revealed that the iridoid pathway in aphids proceeds through the same sequence of intermediates as described for plants. The six identified aphid enzymes are unrelated to their counterparts in plants, conclusively demonstrating an independent evolution of the entire iridoid pathway in plants and insects. In contrast to the plant pathway, at least three of the aphid iridoid enzymes are likely membrane-bound. We demonstrated that a lipid environment facilitates the cyclization of a reactive enol intermediate to the iridoid cyclopentanoid-pyran scaffold in vitro, suggesting that membranes are an essential component of the aphid iridoid pathway. Altogether, our discovery of this complex insect metabolic pathway establishes the genetic and biochemical basis for the formation of iridoid sex pheromones in aphids and this discovery also serves as a foundation for understanding the convergent evolution of complex metabolic pathways between kingdoms.

**Significance Statement:** Plants, animals and microbes produce a plethora of natural products that are important for defense and communication. Most of these compounds show a phylogenetically restricted occurrence, but in rare instances, the same natural product is biosynthesized by organisms in two different kingdoms. The monoterpene-derived iridoids, for example, have been found in more than 50 plant families, but are also observed in several insect orders. The aphid iridoid pathway discovered in this study, one of the longest and most chemically complex insect-derived natural product biosynthetic pathways reported to date, is compared with iridoid biosynthetic pathways in plants and highlights the mechanisms underlying the convergent evolution of metabolic enzymes in insects and plants.

## Introduction

Iridoids are a class of atypical bicyclic monoterpenoids that are widely distributed in flowering plants, but, notably, are also found in several insect orders including Coleoptera, Hymenoptera, and Hemiptera (1). Iridoids therefore present an opportunity to compare and contrast the chemical logic of natural product biosynthesis between plants and insects.

In plants, iridoids largely act as defensive metabolites or biosynthetic intermediates for other natural products (e.g. monoterpenoid indole alkaloids and isoquinoline alkaloids). The pathway leading to the cyclopentanoid-pyran (iridoid) scaffold was first elucidated in the plant Madagascar periwinkle (*Catharanthus roseus*) (2-6) and more recently in the two mint species *Nepeta mussinii* and *N. cataria* (7-9). Iridoid biosynthesis in plants starts with the condensation of the universal terpene precursors isopentenyl diphosphate (IPP) and dimethylallyl diphosphate (DMAPP) to form geranyl diphosphate (GPP), followed by hydrolysis to geraniol (Figure 1A). Both reactions take place in the plastids and are catalyzed by *trans*-isoprenyl diphosphate synthase (IDS) and geraniol synthase (GES), respectively. Hydroxylation of geraniol by geraniol-8-hydroxylase (G8H) leads to 8-hydroxygeraniol, which is further oxidized in two consecutive reaction steps by 8-hydroxygeraniol oxidase (HGO) to 8-oxogeranial. This dialdehyde is then converted to the iridoid nepetalactol by a two-step reduction-cyclization sequence that involves the formation of a highly reactive 8-oxocitronellyl enol/enolate intermediate. Initially, reduction and cyclization of 8-oxogeranial were thought to be controlled by a single enzyme, iridoid synthase (ISY) (3), though later studies showed that ISY likely catalyzes only the NADPH-dependent reduction of 8-oxogeranial to the 8-oxocitronellyl enol/enolate intermediate (8). This intermediate can non-enzymatically cyclize, or, alternatively, the stereoselective cyclization of this intermediate to nepetalactol is enzymatically mediated by nepetalactol-related short-chain dehydrogenase (NEPS) or by major latex protein-like (MLPL) enzymes (8, 9). In *C. roseus*, nepetalactol is further metabolized to secologanin, which serves as a precursor for the formation of monoterpene indole alkaloids in this plant (10). In *Nepeta*, a NEPS protein oxidizes nepetalactol to nepetalactone (8), with both the alcohol and lactone released as volatiles.

**Figure 1:**
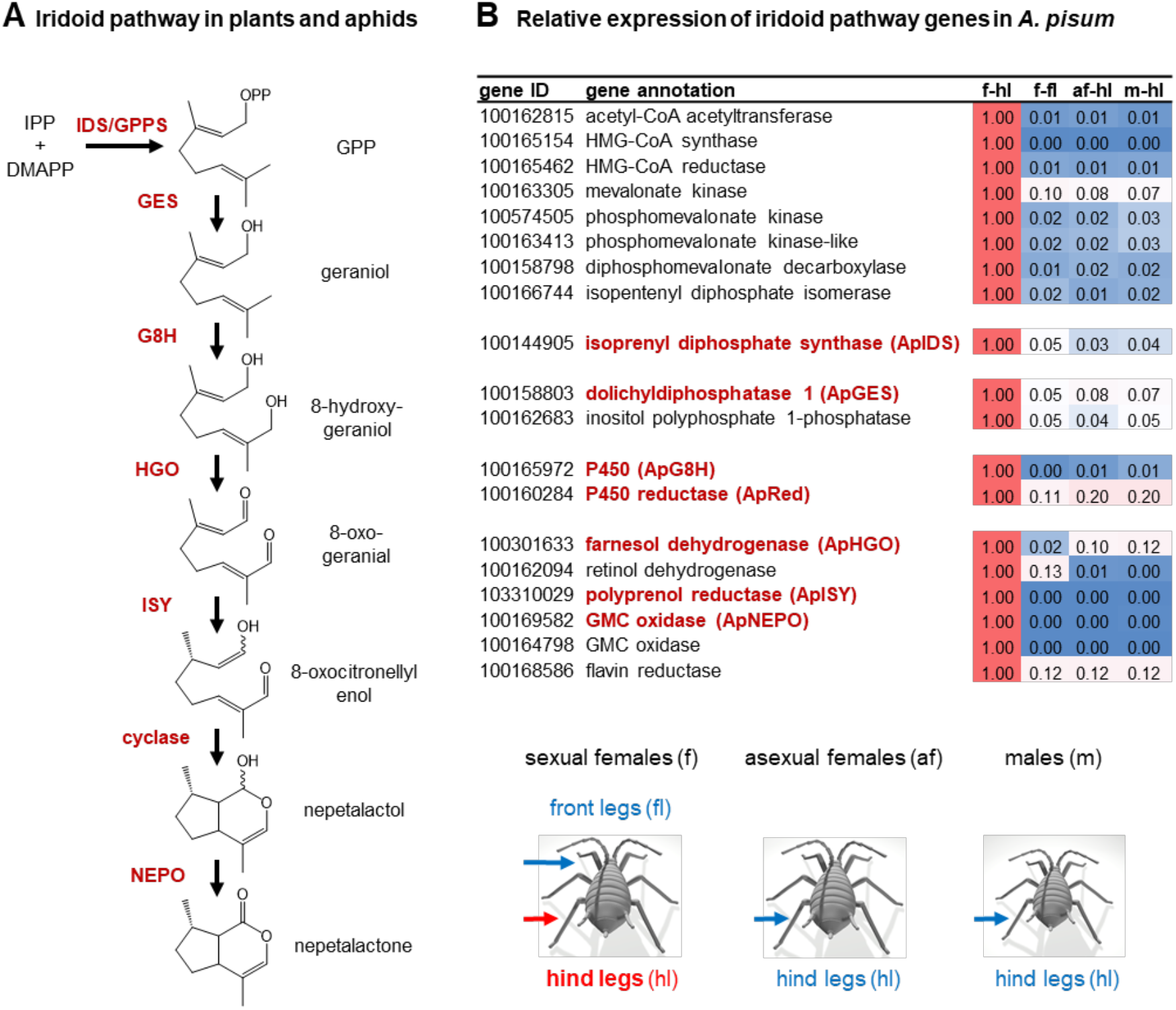
The formation of iridoids in plants and aphids. **(A)** Labeling studies suggest that the biosynthesis of iridoids in the pea aphid *Acyrthosiphon pisum* mimics the biosynthetic pathway in iridoid-producing plants. IPP, isopentenyl diphosphate; DMAPP, dimethylallyl diphosphate; GPP, geranyl diphosphate; IDS, isoprenyl diphosphate synthase; GES, geranyl diphosphate synthase; G8H, geraniol 8-hydroxylase; HGO, 8-hydroxygeraniol oxidoreductase; ISY, iridoid synthase; NEPO, nepetalactol oxidase.**(B)** Relative expression of mevalonate and putative nepetalactone pathway genes in hind legs and front legs of different sexual stages of *A. pisum*. Relative expression data are based on RPKM values obtained by RNAseq. f-hl, hind legs of sexual females; f-fl, front legs of sexual females; af-hl, hind legs of asexual females; m-hl, hind legs of males.

Insects utilize iridoids as both defense compounds and volatile pheromones, but in terms of biosynthesis, comparatively little is understood about insect-derived iridoids. Biosynthetic insights have been obtained from studies on larvae of chrysomelid leaf beetles, which accumulate the iridoid-related monocyclic dialdehydes chrysomelidial and plagiodial (11). Feeding experiments with isotopically-labeled precursors and the discovery of some of the enzymes involved in chrysomelidial formation demonstrated that leaf beetles produce these compounds by a series of chemical reactions similar to those that occur in plants (12-15). Although the enzymatic basis for this pathway has not been completely established, the fact that the known enzymes are unrelated to their counterparts in plants suggests independent evolution of the pathway occurred (14).

*Cis-trans*-nepetalactol and *cis-trans*-nepetalactone are the major iridoids produced by catnip (*N. mussinii*) and catmint (*N. cataria*) (16). These molecules are responsible for the euphoric effect these plants have on cats, but their ecological function is unclear, though they may play roles in mediating interactions with insects (17). Interestingly, *cis-trans*-nepetalactol and *cis-trans*-nepetalactone occur also in aphids, which produce these compounds as volatile sex pheromones (18, 19). The pea aphid *Acyrthosiphon pisum*, for example, has been reported to biosynthesize (1*R*,4*aS*,7*S*,7*aR*)-*cis-trans*-nepetalactol and (4*aS*,7*S*,7*aR*)-*cis-trans*-nepetalactone in glandular structures on the hind legs of sexual female aphids, from where they are released to attract male conspecifics (18, 20). Recent studies with isotopically-labeled iridoid precursors suggest that the iridoid pathway in aphids follows the reaction sequence described for plants (21). However, the underlying enzymatic machinery of this pathway is completely unknown.

Here we report the elucidation of the entire iridoid pathway in the pea aphid *A. pisum*. By searching for genes expressed exclusively in hind legs of sexual female aphids, the site of iridoid production, we could rapidly identify all six biosynthetic genes/enzymes responsible for the conversion of IPP and DMAPP to *cistrans*-nepetalactone. The discovery of the insect nepetalactone pathway in its entirety now allows a comparison of the chemical solutions that have evolved for nepetalactone biosynthesis in plants and animals. Although the chemical steps from GPP to nepetalactone are the same in both *Nepeta* and pea aphids, the enzymes of these pathways have clearly evolved independently.

## Results

### Transcriptome-enabled discovery of iridoid genes in *A. pisum*

Iridoids are produced exclusively in the hind legs of sexual aphid females (20), which allowed us to search for iridoid genes by comparing transcriptomic data from legs of sexual females, asexual females, and males. Since most aphid species have a complex life cycle with multiple asexual generations over the summer and only one sexual female/male generation in fall (22), we subjected a colony of pea aphids to day length/temperature conditions that mimic the fall season to generate sexual females. After verifying that *cis-trans-*nepetalactol and *cis-trans*-nepetalactone were in fact produced in the headspace of the aphids (Figure S1), we then collected hind and front legs of sexual females, hind legs of asexual females, and hind legs of males. RNA was extracted and subjected to Illumnina-sequencing. Out of the 20,918 gene models of the *A. pisum* v3 genome, only 96 appeared to be specifically expressed in hind legs of sexual females (Tables S1 and S2). Notably, among these transcripts were eight genes encoding the entire mevalonate pathway starting from acetyl-CoA to IPP and DMAPP (Figure 1B). Previously reported isotopic labeling studies indicated that the iridoid pathways in pea aphids and plants share at least some of the same biosynthetic intermediates (21). We therefore assumed that the biochemical transformations in pea aphids were the same as those established in *Nepeta*. Then we compiled a list of candidates from the 96 genes specifically expressed in sexual female hind legs that also encoded enzymes that could, in principle, carry out these predicted reactions. One putative *IDS* gene, two putative phosphatase genes, one putative cytochrome P450 gene, one putative P450 reductase gene, and six putative oxidase/reductase genes were selected for further characterization (Figure 1B).

### ApIDS catalyzes the metal ion cofactor-dependent formation of GPP

The formation of GPP in the horseradish leaf beetle *Phaedon cochleariae* is catalyzed by a bifunctional IDS (PcIDS) that shows a metal ion cofactor-dependent product specificity, producing primarily GPP with cobalt or manganese, or farnesyl diphosphate (FPP) with magnesium (12). A homologue of PcIDS (ApIDS, gene ID, 100144905) was specifically expressed in sexual female hind legs. Phylogenetic analysis revealed that PcISD and ApIDS clustered together in a clade of beetle and aphid GPP/FPP synthases (Figure 2A). In vitro characterization of recombinant ApIDS (Tables S3 and S4) showed IDS activity when incubated with IPP and DMAPP. Using magnesium as a cofactor, ApIDS produced similar amounts of GPP, FPP, and geranylgeranyldiphosphate (GGPP), while cobalt and, to a lesser extent, manganese shifted the product specificity to GPP as the major product (Figure 2B, Figure S2). This suggests that ApIDS, like PcIDS, uses cobalt or manganese as metal ion cofactor to produce GPP for iridoid formation in vivo.

**Figure 2:**
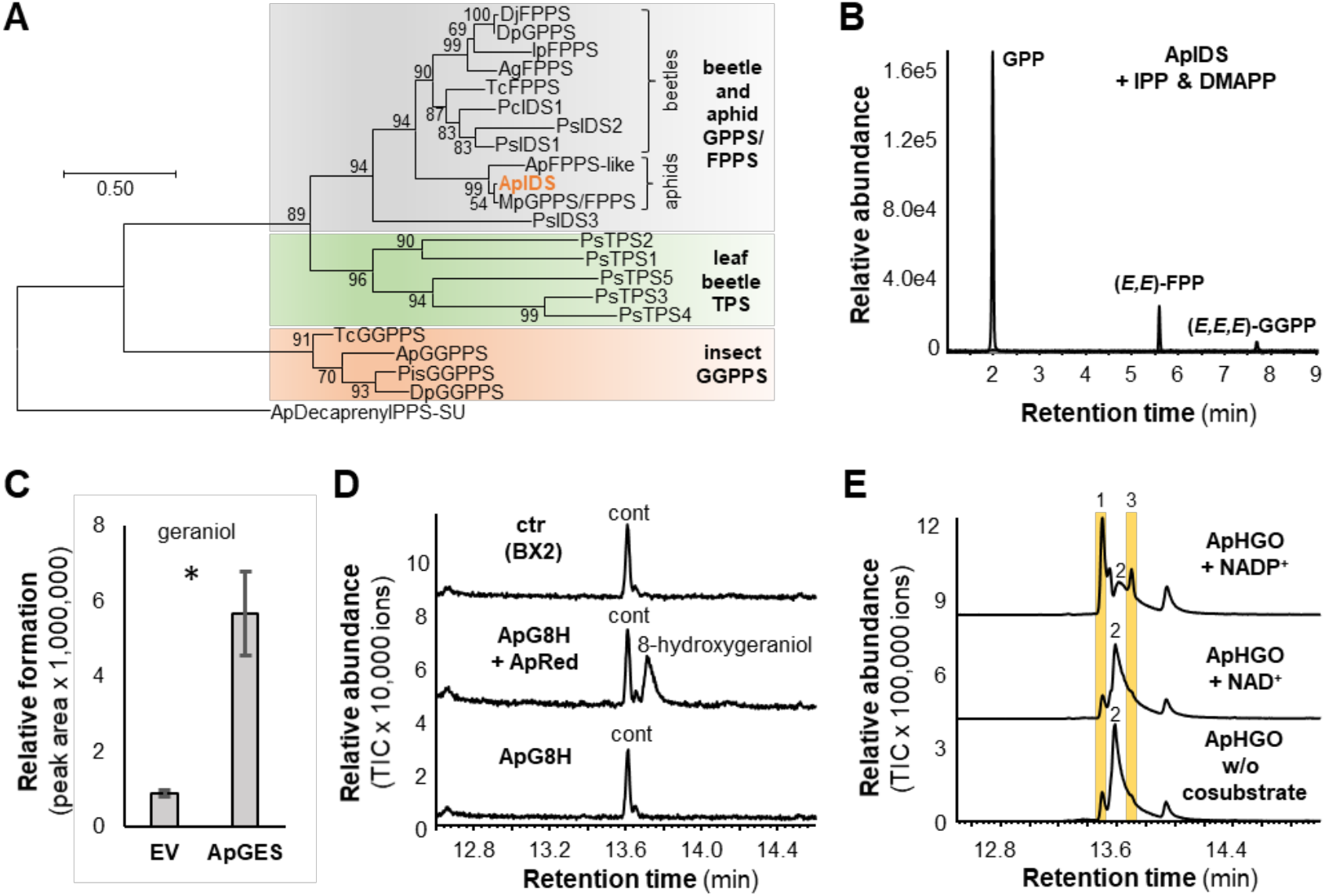
Biochemical characterization of ApIDS, ApGES, ApG8H, and ApHGO. **(A)** Cladogram analysis of characterized isoprenyldiphosphate synthases (IDS) and terpene synthases (TPS) from insects. The tree was inferred by using the Maximum Likelihood method based on the JTT model. Bootstrap values (n = 1000) are shown next to each node. The rooted tree is drawn to scale, with branch length measured in the number of amino acid substitutions per site. Putative decaprenyldiphosphate synthase subunit 2 from *Acyrthosiphon pisum* was used as an outgroup. **(B)** Characterization of ApIDS. ApIDS lacking the N-terminal signal peptide was expressed as an N-terminal His-tag-fusion protein in *Escherichia coli*, purified, and incubated with the substrates IPP and DMAPP in the presence of 1 mM CoCl_2_. Enzyme products GPP, (*E,E*)-FPP, and (*E,E,E*)-GGPP were analyzed using liquid chromatography-tandem mass spectrometry. **(C)** Characterization of ApGES. The phosphatase ApGES was expressed in *Saccharomyces cerevisiae* and microsomes harboring the recombinant protein were incubated with GPP. Geraniol was extracted with hexane and analyzed using gas chromatography-mass spectrometry (GC-MS). Mean peak area ± SE are shown (n = 3 independent expression constructs). The amount of geraniol formed was significantly higher in ApGES samples than in the negative controls (empty vector (EV)) (t = 5.219; df = 2.026; p = 0.034). **(D)** Characterization of ApG8H. *S. cerevisiae* microsomes containing either ApG8H, ApG8H in combination with the P450 reductase ApRed, or the maize P450 BX2 as negative control were assayed with geraniol as substrate and NADPH as cosubstrate. Reaction products were analyzed using GC-MS. cont, di-tert-butylphenol (contamination). **(E)** Characterization of ApHGO. ApHGO was expressed as N-terminal His-tag-fusion protein in *E. coli*, purified, and incubated with 8-hydroxygeraniol either in the absence or presence of NAD(P). Enzyme products were extracted from the assays and analyzed using GC-MS. 1, 8-oxogeraniol (partially oxidized product); 2, 8-hydroxygeraniol (starting material); 3, 8-oxogeranial (fully oxidized product). Note: The tailing of the peaks is due to the polar nature of the aldehydes and the alcohol.

### The phosphatase ApGES hydrolyzes GPP to geraniol

In plants, the formation of the mono- and sesquiterpene alcohols geraniol and farnesol from GPP and FPP is catalyzed by terpene synthases (TPS) (23). However, in insects, farnesol, which is an intermediate in juvenile hormone biosynthesis, is produced from FPP by a phosphatase belonging to the haloalkanoic acid dehalogenase (HAD) super family (24, 25). Our candidate search in the pea aphid transcriptome did not reveal any TPS-or HAD-like proteins that were selectively expressed in sexual female hind leg. Instead, we identified two putative phosphatases annotated as dolichyldiphosphatase (gene ID, 100158803) and inositol polyphosphate-1-phosphatase (gene ID, 100162683) (Figure 1B) (Tables S3 and S5). The putative inositol polyphosphate-1-phosphatase, expressed in *Escherichia coli* as a soluble protein, showed no GPP hydrolysis activity. In contrast, *Saccharomyces cerevisiae* microsomes harboring this recombinant membrane-bound putative dolichyldiphosphatase protein hydrolyzed GPP to geraniol (Figure 2C, Figure S3). Thus, we named this dolichyldiphosphatase homologue ApGES.

### The P450 ApG8H and the P450 reductase ApRed act together to produce 8-hydroxygeraniol

Both, plants and the leaf beetle *P. cochleariae* utilize P450 enzymes to catalyze the hydroxylation of geraniol to 8-hydroxygeraniol (2, 9, 15). Only a single P450 in the pea aphid transcriptome displayed selective expression in sexual female hind legs, and this enzyme grouped together with PcG8H from *P. cochleariae* in a phylogenetic analysis (Figure 1B, Figure S4), though these two proteins share only 35 % amino acid sequence identity. The aphid gene was named *ApG8H* and the complete ORF was expressed in *S. cerevisiae* either alone or together with a putative P450 reductase gene (*ApRed*; gene ID, 100162683), which had a similar expression pattern as *ApG8H*. In the presence of the cosubstrate NADPH and ApRed, ApG8H converted geraniol to 8-hydroxygeraniol (Figure 2D). A heterologously expressed P450 from maize (BX2 (26)) that was used as negative control, and ApG8H expressed without ApRed, showed no activity. Notably, ApG8H exhibited a relatively broad substrate specificity, hydroxylating citronellol, nerol, linalool, and neral, though not the monoterpene hydrocarbons limonene and myrcene (Figure S5), suggesting that an oxygen atom at C1 is critical for binding or catalysis.

### The short-chain reductase ApHGO catalyzes the NADP^+^-dependent oxidation of 8-hydroxygeraniol to the iridoid precursor 8-oxogeranial

While the oxidation of 8-hydroxygeraniol to 8-oxogeranial in plants is catalyzed by alcohol dehydrogenases (5, 9), *P. cochleariae* beetles use a flavin-dependent glucose-methanol-cholin (GMC) oxidase to catalyze this reaction (13). Thus, we tested two aphid short chain alcohol dehydrogenase (SDR) candidates, one annotated as farnesol dehydrogenase (gene ID, 100301633) and the other as retinol dehydrogenase (gene ID, 100162094), as well as two GMC candidates (gene IDs, 100169582 and 100164798) that were selectively expressed in sexual female hind leg (Figure 1B; Figure S6). The complete ORFs were expressed in *E. coli* and purified proteins were assayed with 8-hydroxygeraniol in the presence of NADP^+^. Enzyme activity could only be observed for the putative farnesol dehydrogenase (named ApHGO), which catalyzed this two-step oxidation (Figure 2E, Figure S7). Further characterization showed that ApHGO preferred NADP^+^ over NAD^+^ as cosubstrate and exhibited a broader substrate specificity, also oxidizing geraniol and nerol, but not β-citronellol (Figure S8). The putative retinol dehydrogenase 100162094 and the GMC oxidase 100169582, although not able to accept 8-hydroxygeraniol as substrate, converted geraniol to geranial (Figure S7).

### The iridoid synthase ApISY is a membrane protein catalyzing the reduction of 8-oxogeranial

The crucial formation of the cyclopentanoid-pyran scaffold occurs with the reductive cyclization of 8-oxogeranial. In plants, this step is initiated by iridoid synthase, an SDR that belongs to the progesterone 5β-reductase super family (3). The enzyme that insects use to catalyze this reduction is unknown. In vitro assays of a putative retinol dehydrogenase (gene ID, 100162094) and two GMC oxidases (gene IDs, 100169582 and 100164798) from our list of candidate genes showed no activity with the substrate 8-oxogeranial and the cosubstrate NADPH (Figure S9). However, a putative oxidoreductase annotated as membrane-bound polyprenol reductase (gene ID, 103310029 (Table S5)) was also selectively expressed in sexual female hind legs. Yeast microsomes containing this recombinant protein converted the substrate 8-oxogeranial to *cis-trans*-nepetalactol as the major product, along with minor amounts of *cis-trans*-iridodial and five other unidentified compounds (Figure 3A; Figure S9; Figure S10). Thus, the tested protein was named ApISY. As with plant iridoid synthases, without NADPH, ApISY showed no activity (Figure 3A).

**Figure 3:**
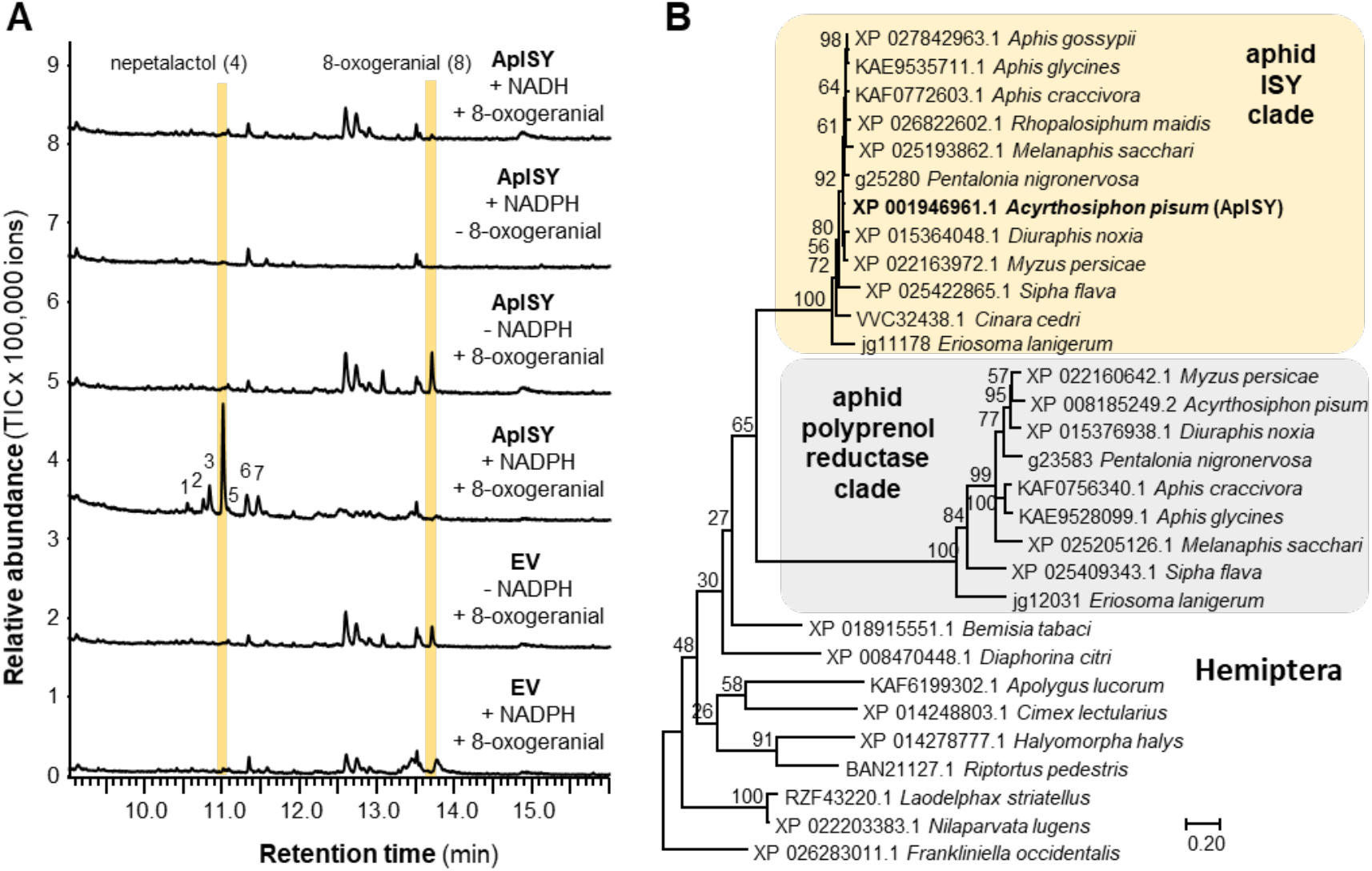
ApISY is a membrane-bound reductase that likely evolved from a polyprenol reductase ancestor. **(A)** Yeast (*Saccharomyces cerevisiae*) microsomes made from a strain harboring the empty expression vector (EV) or expressing ApISY (ApISY) were assayed with 8-oxogeranial either in the presence or absence of NAD(P)H. Reaction products were extracted with ethyl acetate and analyzed using GC-MS. **(B)** Rooted tree of polyprenol reductases and iridoid synthases from Hemiptera. The tree was inferred by using the Maximum Likelihood method based on the Le_Gascuel_2008 model. Bootstrap values (n = 1000) are shown next to the branches. A discrete Gamma distribution was used to model evolutionary rate differences among sites. The tree is drawn to scale, with branch lengths measured in the number of amino acid substitutions per site. All positions with less than 95% site coverage were eliminated. A putative polyprenyol reductase from the Western flower thrips, *Frankliniella occidentalis*, was used to root the tree. NCBI accession numbers for all sequences are given in the tree.

In *Nepeta*, and likely also in other plants, iridoid synthase works in concert with cyclases that mediate the stereoselective cyclization of the initial reduction product of ISY, 8-oxocitronellyl enol/enolate, into different nepetalactol stereoisomers (8). When any plant ISY is assayed in vitro without a cyclase, the product profile is strongly dependent on the assay conditions. In high buffer concentrations or at low pH values, spontaneous tautomerization of the 8-oxocitronellyl enol/enolate intermediate to 8-oxocitronellal is favored, while low buffer conditions or higher pH values lead to the spontaneous cyclization to *cis-trans*-nepetalactol as the predominant product. Moderate buffer concentrations or pH values lead to a mixture of monocyclic dialdehydes (8). Using ISY from the plant *C. roseus* (CrISY) as a point of comparison (Figure 4; Figure S11), we were surprised to observe that assays with yeast microsomes containing ApISY showed specificity for formation of *cis-trans*-nepetalactol over a broad buffer concentration range. Only at a buffer concentration of 0.5 M, yield of *cis-trans*-nepetalactol was affected in the ApISY reaction (Figure 4). Since the plant ISY is a soluble protein, while ApISY is an integral membrane protein with seven predicted transmembrane domains (Table S5), we assayed CrISY in the presence of yeast microsomes to determine whether the membranes had an impact on product profile. Notably, *cis-trans*-nepetalactol was the predominant product of CrISY throughout the buffer concentration range after addition of microsomes (Figure 4). This suggests that the lipid environment of the membranes likely facilitates spontaneous cyclization to the iridoid scaffold by preventing contact of the reactive intermediate with general acid catalysts such as buffer components in the assay.

**Figure 4:**
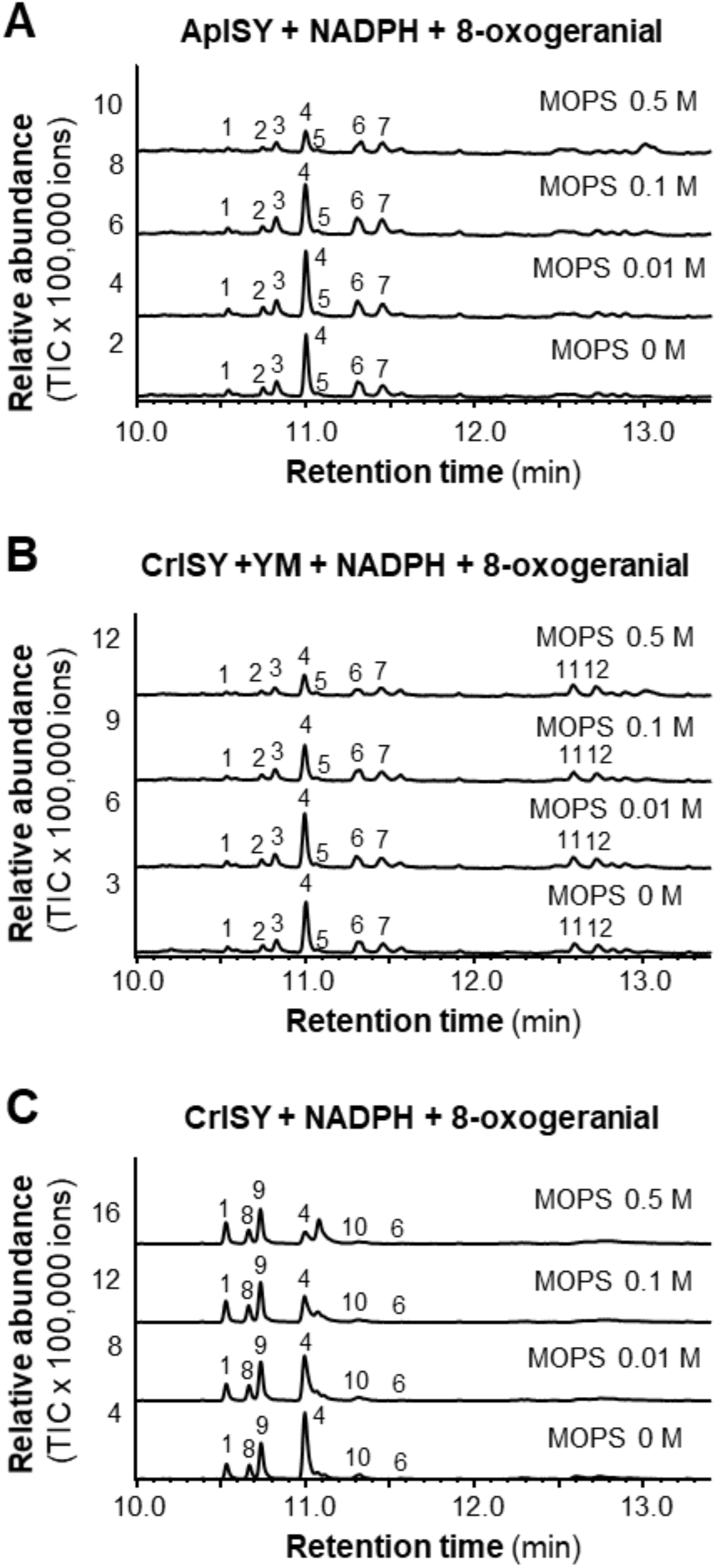
Buffer concentration and the lipid environment influence iridoid synthase activity. **(A)** Yeast (*Saccharomyces cerevisiae*) microsomes containing ApISY were assayed with 8-oxogeranial in the presence of NADPH under different buffer concentrations. **(B)** CrISY was expressed in *Escherichia coli*, purified, and assayed with 8-oxogeranial and NADPH under different buffer concentrations in the background of yeast microsomes (YM). **(C)** Purified CrISY was incubated with 8-oxogeranial and NADPH under different buffer concentrations. Reaction products were extracted with ethyl acetate and analyzed using gas chromatography-mass spectrometry. 1, *cis-trans*-iridodial; 2, unidentified; 3, unidentified; 4, *cis-trans*-nepetalactol; 5, unidentified; 6, unidentified; 7, unidentified; 8, *trans-trans*-iridodial; 9, *cis-trans*-iridodial; 10, tetrahydro-8-oxogeranial; 11, unidentified; 12, unidentified.

### ApISY likely evolved from a polyprenol reductase ancestor

Polyprenol reductases, ubiquitous in Eukaryotes, catalyze reduction of the α-isoprene unit of polyprenols to form dolichols, the precursors for dolichol-linked monosaccharides that are required for protein N-glycosylation (27-29). Together with steroid 5α-reductases and very-long-chain enoyl-CoA reductases, polyprenol reductases belong to the steroid 5α-reductase (SRD5A) family (Pfam, PF02544). A BLAST analysis with ApISY as query revealed two polyprenol reductase-like genes in most of the available aphid genomes. In a phylogenetic analysis, these genes formed two distinct and aphid-specific clades among the polyprenol reductases of eukaryotes (Figure 3B; Figure S12).

### The flavin-dependent GMC oxidase ApNEPO converts *cis-trans*-nepetalactol into *cis-trans*-nepetalactone

The SDR NEPS1 from *Nepeta* catalyzes the oxidation of *cis-trans*-nepetalactol to *cis-trans*-nepetalactone

(8). To identify the enzyme that catalyzes this reaction in pea aphids, the putative retinol dehydrogenase (gene ID, 100162094) and the two FAD-dependent GMC oxidases (gene IDs, 100169582 and 100164798) from our candidate gene list (Figure 1B) were assayed with the *cis-trans*-nepetalactol substrate and NADP^+^. While the SDR 100162094 and the GMC oxidase 100164798 were not active, GMC oxidase 100169582 converted *cis-trans*-nepetalactol into *cis-trans*-nepetalactone (Figure 5, Figure S13). A phylogenetic analysis revealed that this enzyme, designated *A. pisum* nepetalactol oxidase (ApNEPO), belonged to the ε clade of GMC oxidoreductases (Figure S14) and was not related to the leaf beetle enzyme PcHGO, which clustered into a beetle-specific GMC clade (Figure S14). Sequence prediction suggested that ApNEPO contains a signal peptide targeting the protein into the lumen of the endoplasmatic reticulum (Table S5).

**Figure 5:**
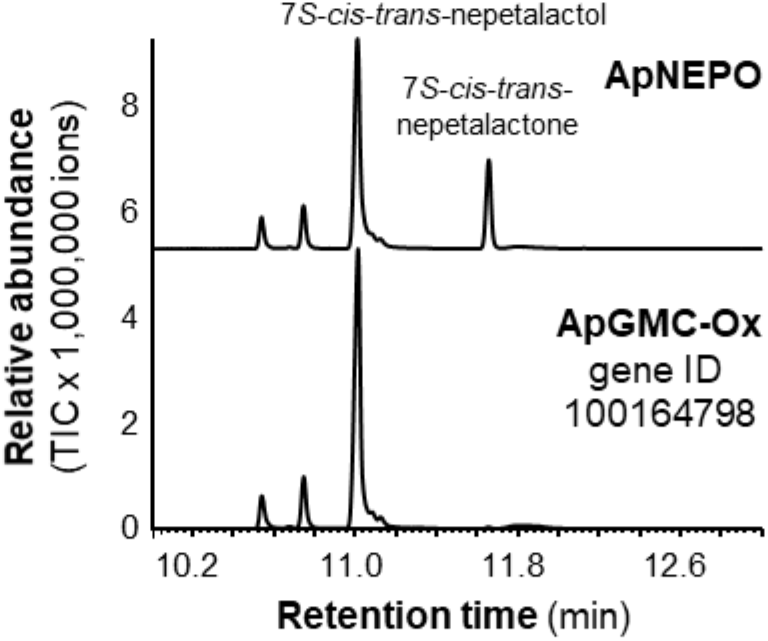
Biochemical characterization of ApNEPO. ApNEPO and another putative GMC-oxidase (ApGMC-Ox) highly expressed in hind legs of *A. pisum* sexual females were expressed as N-terminal His-tag-fusion proteins in *Escherichia coli*, purified, and incubated with 7*S*-*cis*-*trans*-nepetalactol in the presence of NADP. Enzyme products were extracted with ethyl acetate and analyzed using gas chromatography-mass spectrometry.

## Discussion

Here we elucidated the entire iridoid pathway in the pea aphid *A. pisum*. Previously reported feeding studies in pea aphids (21), as well as the spatial localization of nepetalactone biosynthesis (20), guided the identification of the six biosynthetic genes (Figure 1A). The characterization of these enzymes indicates that the plant and aphid nepetalactone biosynthetic pathways are composed of the same chemical transformations. However, each of the respective enzymes clearly evolved independently in plants and aphids. Our data show that in some cases plants and aphids recruited enzymes from different protein families to catalyze the same reactions (Figure S15). The formation of geraniol in *Catharanthus* and *Nepeta*, for example, is mediated by terpene synthases (5, 9), while the pea aphid uses a phosphatase to produce geraniol by direct hydrolysis of the phosphodiester bond of GPP (Figure 2C). Moreover, the reductive cyclization of 8-oxogeranial to *cis-trans*-nepetalactol and the subsequent oxidation of this alcohol to *cistrans*-nepetalactone in aphids involves the action of a polyprenol reductase-like protein and a flavin-dependent oxidase from the GMC family, respectively, while plants recruited members of the SDR family to catalyze both reactions (Figure S15).

Only one other natural product pathway, the three-step biosynthesis of the cyanogenic glycoside linamarin, has been fully elucidated in both plants and insects. Recent work has demonstrated that linamarin biosynthesis consists of two cytochrome P450s and one glucosyl transferase in both plants and insects, and that these pathways arose independently (30, 31). Additionally, terpene synthases, the key enzymes in terpene formation, have been identified in both plants and insects, and these enzymes are also the result of independent evolution in the different kingdoms (32-34).

Iridoids and iridoid-related compounds are widespread among insects and have been observed in different insect orders including Coleoptera, Hymenoptera, and Hemiptera (1). The discovery of the aphid nepetalactone pathway provides an opportunity to determine whether iridoids evolved convergently in two divergent species of insects. The biosynthesis of the iridoid-related dialdehyde chrysomelidial has been partially elucidated in the leaf beetle *P. cochleariae* (11). Although chrysomelidial lacks the cyclopentanoid-pyran scaffold that defines the iridoids, its formation shares many of the same reactions as the iridoid core pathway in aphids (Figure S15). GPP is produced in the leaf beetle by PcIDS, which is obviously phylogenetically related to ApIDS (12) (Figure 2A). Geraniol, which is produced in beetles by an as yet undiscovered enzyme, acts as substrate for PcG8H, a P450 that hydroxylates this alcohol to 8-hydroxygeraniol (15). Although PcG8H and ApG8H both belong to clan 3 of insect P450s (Figure S4), their low sequence identity suggests independent origins. Independent evolution of enzyme activities is even more obvious for the last two steps of the chrysomelidial pathway. The oxidation of 8-hydroxygeraniol in *P. cochleariae* is catalyzed by a GMC oxidase (13), in contrast to aphids, which recruited an SDR for the same reaction, and the final non-reductive cyclization of the formed 8-oxogeranial to chrysomelidial is presumably mediated by a PcTo-like juvenile hormone-binding protein, although conclusive evidence of this enzyme activity is still lacking (35). Overall, the elucidation of the iridoid pathway in aphids presented here shows that although the reaction sequence is conserved, iridoid formation evolved independently not only in different kingdoms, but also in different insect orders through convergent evolution (Figure S15).

From a chemical perspective, the first committed step in the iridoid pathway, the cyclization of 8-oxogeranial by iridoid synthase (ISY), is of mechanistic interest. All known plant iridoid synthases belong to the SDR protein family and have been described to catalyze the reduction of 8-oxogeranial to a highly reactive 8-oxocitronellyl enol/enolate intermediate, which is then cyclized by NEPS or MLPL proteins to different stereoisomers of nepetalactol (8, 9). In the absence of a cyclase, 8-oxocitronellyl enol/enolate can react spontaneously to form various compounds depending on the assay conditions. Higher pH values or low buffer concentrations lead to cyclization to *cis-trans*-nepetalatol, while acidic conditions or high buffer concentrations favor spontaneous formation of dialdehydes or 8-oxocitronellal (8). In contrast to plant ISY, a soluble protein likely located in the cytosol, the iridoid synthase in the pea aphid is an integral membrane protein predicted to possess seven membrane domains (Table S5). Interestingly, yeast microsomes harboring ApISY produced mainly *cis-trans*-nepetalactol independent of the buffer concentration (Figure 4A). Moreover, when CrISY from the plant *C. roseus* was tested in the presence of microsomes prepared from a control yeast strain, the same trend, the production of mainly *cis-trans*-nepetalactol at all buffer concentrations tested, was observed (Figure 4B). This indicates that the lipid environment of membranes favors the spontaneous cyclization of the 8-oxocitronellyl enol/enolate to *cis-trans*-nepetalactol, presumably by preventing contact of the reactive intermediate with protons or other acidic compounds. Thus, in principle, iridoid formation in pea aphids does not require the action of a cyclase. However, we cannot rule out that ApISY itself or other, as yet unidentified, aphid proteins fulfill this function in vivo. Most aphid species produce *cis-trans*-nepetalactol (18, 36). The damson-hop aphid *Phorodon humili*, however, produces the *cis-cis* isomer (37, 38). This species must have either an iridoid synthase with different stereoselectivity, catalyzing both the reduction and cyclization of 8-oxogeranial to the final nepetalactol stereosiomer, or a partner cyclase that cyclizes the potential 8-oxocitronellyl enol/enolate intermediate to *cis-cis*-nepetalactol. Elucidating the biosynthetic pathway for *cis-cis*-nepetalactol in *P. humili* would provide additional insight into the complex chemistry underlying the formation of the iridoid backbone in animals.

Sequence comparisons revealed that ApISY is related to polyprenol reductases, a class of integral membrane proteins involved in N-glycosylation of secreted and membrane-bound proteins (27-29) (Figure S12). Interestingly, most of the aphid species sequenced to date possess two putative polyprenol reductase copies that form two distinct and aphid-specific clades in a phylogenetic tree of the polyprenol reductases of Hemiptera (Figure 3B; Figure S12). Thus, it is likely that the ApISY-containing clade represents aphid iridoid synthases, while the other clade contains true polyprenol reductases, which is a metabolic enzyme that is essential for survival. The close relationship of these two clades suggests that iridoid synthase activity arose by gene duplication and subsequent neofunctionalization of a polyprenol reductase gene early in aphid evolution or in an ancestor of the aphids. Furthermore, the striking sequence similarities among proteins within the two clades (Figure 3B) indicates a high degree of purifying selection to preserve their respective enzymatic functions. We cannot predict the evolutionary origin of other iridoid pathway genes because the function of their closest homologues in aphids are still unknown. For example, although ApGES was annotated as a dolichyldiphosphatase, an enzyme acting together with polyprenol reductase in the process of N-glycosylation of secreted and membrane-bound proteins (39, 40), there is no experimental evidence for dolichyldiphosphatase activity of ApGES, and BLAST analysis did not reveal any other *ApGES*-related gene that could be dedicated to dolichyldiphosphatase function in the *A. pisum* genome.

A defining feature of the iridoid biosynthetic pathway in aphids is that many of the enzymes appear to be membrane anchored. In addition to the iridoid synthase ApISY, the phosphatase ApGES, the cytochrome P450 ApG8H and its reductase ApRed were predicted to be integral membrane proteins (Table S5). Moreover, the prediction of an endoplasmic reticulum (ER) signal peptide in ApNEPO suggests that this protein is localized in the lumen of the ER (Table S5). This leads us to speculate that these enzymes may form a metabolon, most likely, based on signal sequence prediction, on the membrane of the ER (Figure 6). Given the predicted mitochondrial signal peptide of ApIDS, the pathway likely starts in the mitochondria with the formation of GPP, which is then transported to the ER. Notably, the list of candidate iridoid genes exclusively expressed in hind legs of sexual female aphids contained seven genes annotated as transporters for the inner and outer membrane of mitochondria that could be involved in GPP transport (Table S1). The hydrolysis of GPP and the formation of the iridoid backbone could then be catalyzed by the putative metabolon in the ER membrane, which may provide efficient substrate channeling, preventing the release of highly reactive pathway intermediates such as 8-oxogeranial. The final oxidation of the comparatively stable nepetalactol product to nepetalactone could then occur in the lumen of the ER, and formation of nepetalactone-containing vesicles and their transport to the cell membrane might represent a possible mechanism for the active release of these volatile iridoids.

**Figure 6:**
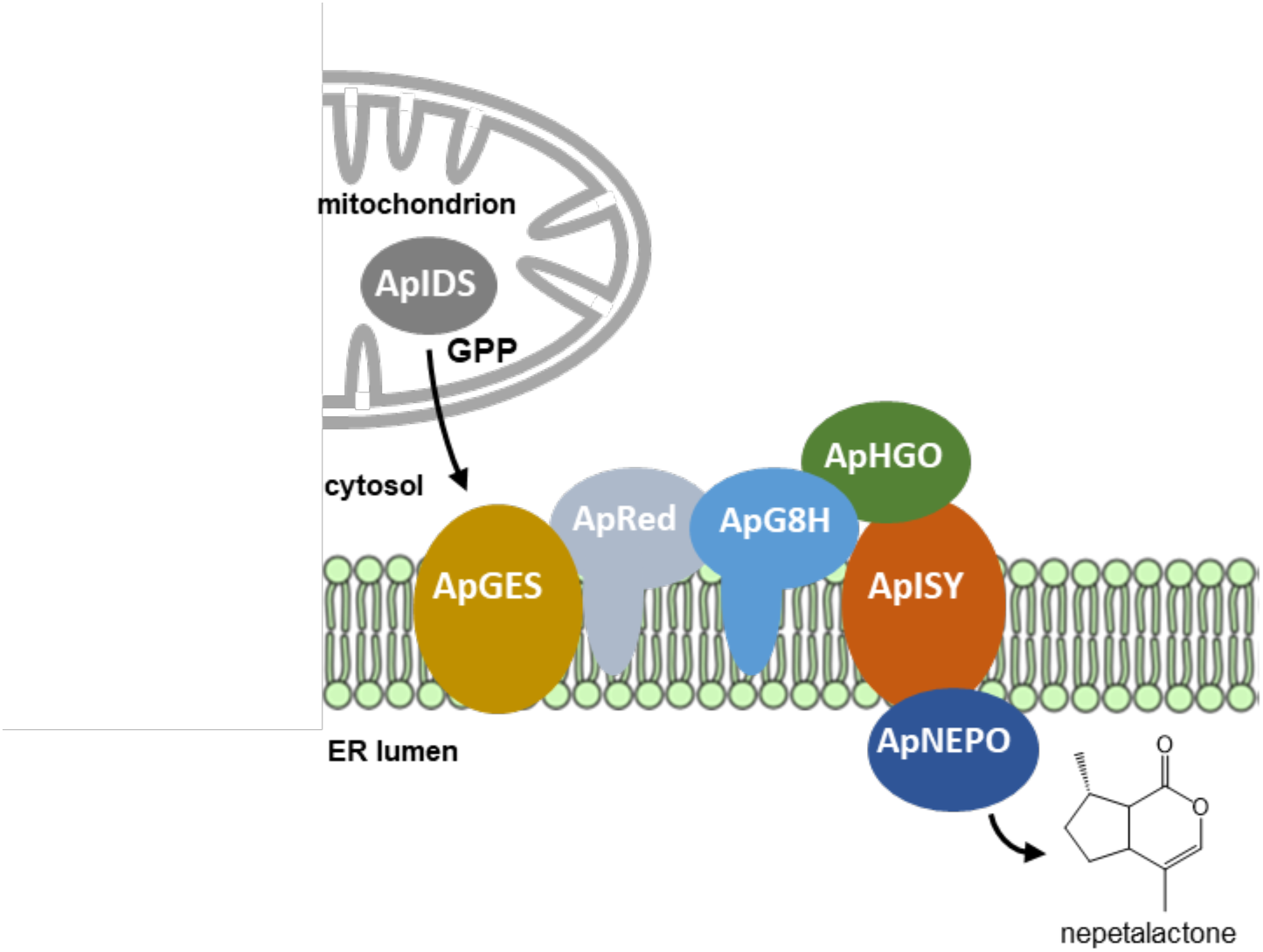
The iridoid pathway in aphids might be organized in a membrane-associated metabolon. While ApHGO is predicted to be localized as soluble protein in the cytosol and ApNEPO is likely localized in the lumen of the endoplasmic reticulum (ER), ApGES and ApISY are transmembrane proteins.

Overall, chemical logic, along with the discreet spatial localization of the site of biosynthesis, facilitated the discovery of the six-step pathway for nepetalactone biosynthesis in animals. This provides a foundation for understanding how complex natural products have evolved in two kingdoms of life. The insect pathway also provides new insights into the relatively understudied field of insect natural product biosynthesis.

## Materials and Methods

### Cultivation of *Acyrthosiphon pisum* and generation of sexual female aphids

Asexual females of pea aphid (*Acyrthosiphon pisum* Harris) clone JML06 were reared on 4 week-old broad bean (*Vicia faba*) cv. “The Sutton” plants under long-day conditions (16/8 h light/dark, 22°C, 60 % humidity). To avoid escape of aphids, plants were covered with air-permeable cellophane bags (18 × 38.5 cm, Griesinger Verpackungsmittel, Neuenbürg - Arnbach, Germany). To generate sexual female and male aphids, asexual L3 aphid larvae were transferred to short-day conditions mimicking fall season (12/12 h light/dark, 14°C, 60 % humidity). Two generations later, sexual females and males were produced, and adult aphids (6-10 day old) were used for experiments. The emission of iridoids by sexual female aphids was tested by placing a solid phase microextraction (SPME) fiber for three hours into the headspace of *V. faba* plants with the aphids. The SPME fiber was then loaded into the injector of a gas chromatograph coupled with a mass spectrometer as described below.

### Transcriptome sequencing and gene identification

For RNA extraction, 20 hind legs of sexual females, 20 front legs of sexual females, 20 hind legs of male aphids, and 20 hind legs of asexual aphids were collected and directly placed in 450 µl lysis buffer containing guanidinium thiocyanate (innuPREP RNA Mini Kit, IST Innuscreen, Berlin, Germany). Material was shredded by shaking with metal beads using a Tissue Lyzer II (Qiagen) for 2 × 4 min (frequency 50/s). Total RNA was extracted with the innuPREP RNA Mini Kit according to manufacturer’s instructions, eluted in 30 µl RNAse free water, and sent to Novogene (Cambridge, UK) for RNA-seq library construction (polyA enrichment) and sequencing (NovaSeq PE150, paired reads, 6 gigabyte of raw data per sample). Trimming of the obtained sequencing reads and mapping to the pea aphid genome (version 3) were performed with the program CLC Genomics Workbench (Qiagen Bioinformatics) (mapping parameter: length fraction, 0.8; similarity fraction, 0.9; max number of hits, 25). Raw reads were deposited in the NCBI Sequence Read Archive (SRA) under the BioProject accession PRJNAxxx. In order to identify pea aphid genes involved in iridoid formation, we performed Pearson correlation based on the hypothesis that iridoid genes are exclusively expressed in hind legs of sexual female aphids. Genes with a Pearson correlation coefficient ≥ 0.99, a RPKM value ≥ 10 in hind legs of sexual female aphids, and a fold change ≥ 5 (hind legs of sexual female aphids versus other samples) were considered as candidates (Table S1).

### Prediction of signal peptides and transmembrane domains

Prediction of signal peptides (Table S3) and transmembrane domains (Table S5) was performed using TargetP-2.0 (https://services.healthtech.dtu.dk/service.php?TargetP-2.0) and DeepTMHMM (https://dtu.biolib.com/DeepTMHMM/), respectively.

### Gene Synthesis and cloning

The complete open reading frames (ORF) of *ApHGO, ApNEPO*, 100162094, 100164798, 100162683, and 100168586, as well as the N-terminal truncated ORF of *ApIDS* lacking the predicted signal peptide were synthesized after codon optimization for heterologous expression in *E. coli* by Twist Bioscience and inserted as *Bam*HI/*Hin*dIII fragments into the vector pET-28a(+) that allows expression as N-terminal His-tag fusion protein in *E. coli. ApGES* was codon-optimized for *S. cerevisiae* and synthesized by Twist Bioscience. The complete ORFs of *ApG8H, ApRed*, and *ApISY* were amplified from cDNA obtained from hind legs of sexual female pea aphids using the primers listed in Table S6. *ApG8H* and *ApRed* were cloned as sticky-end fragments into the same pESC-Leu 2d vector using the two different cloning sites (41). *ApISY* and *ApGES* were separately cloned as sticky-end fragments into pESC-Leu 2d. cDNA was synthesized from total RNA (1 µg) treated with DNaseI (Thermo Fisher Scientific) using SuperScript III reverse transcriptase and oligo (dT)20 primers (Invitrogen) according to the manufacturer’s instructions. All synthesized or amplified sequences are given in Table S4. Amplified sequences were deposited in NCBI GenBank with the accession numbers ON862918 (*ApG8H*), ON862919 (*ApRed*), and ON862920 (*ApISY*).

### Heterologous expression of candidate genes in *Escherichia coli*

Expression constructs were transferred to *E. coli* strain BL21 (DE3) (Invitrogen). Liquid cultures were grown in lysogeny broth at 37°C and 220 rpm until an OD600 of 0.7, induced with a final concentration of 0.5 mM IPTG, and subsequently incubated at 18°C and 220 rpm for 16 h. The cells were harvested by centrifugation at 3200 *g* for 10 min, resuspended in refrigerated extraction buffer (50 mM Tris-HCl pH 8, 500 mM NaCl, 20 mM imidazole, 5% (v/v) glycerol, 50 mM glycine, EDTA-free protease inhibitor (1 tablet/50 mL buffer, freshly added), and lysozyme (10 mg/50 mL buffer, freshly added)) and disrupted by sonication for 2 min (2s on, 3s off) on ice (Bandelin UW 2070). Cell debris were removed by centrifugation (35,000 *g* at 4°C for 20 min) and the N-terminal His-tagged proteins were purified from the supernatant using NiNTA agarose (Qiagen) according to the manufacturer’s instructions. The buffer of the eluted protein samples was exchanged for assay buffer (for details see paragraph enzyme assays) 100 mM MOPS pH 7.5, 10% (v/v) glycerol) by using Amicon 10K Concentrator columns (Merck Millipore). SDS-polyacrylamid gel electrophoresis and spectrophotometric analysis was used to check purity and approximate quantity of proteins.

### Heterologous expression of ApGES, ApG8H, and ApISY in *Saccharomyces cerevisiae*

For heterologous expression in yeast, constructs were transformed into the *S. cerevisiae* strain INVSc1 (Thermo Fisher) using the S.c. EasyComp Transformation Kit (Invitrogen) according to the manufacturer’s instructions. Subsequently, 30 mL Sc-Leu minimal medium (6.7 g/L yeast nitrogen base without amino acids, but with ammonium sulfate; 100 mg/L of each L-adenine, L-arginine, L-cysteine, L-lysine, L-threonine, L-tryptophan, and uracil; 50 mg/L of each L-aspartic acid, L-histidine, L-isoleucine, L-methionine, L-phenylalanine, L-proline, L-serine, L-tyrosine, L-valine; 20 g/L d-glucose) was inoculated with single yeast colonies and grown overnight at 28°C and 180 rpm. For main cultures, 100 mL YPGA (Glc) full medium (10 g/L yeast extract, 20 g/L bactopeptone, 74 mg/L adenine hemisulfate, 20 g/L d-glucose) was inoculated with one unit OD600 of the overnight cultures and incubated under the same conditions for 30-35 h. After centrifugation (5000 *g*, 16°C, 5min), the expression was induced by resuspension of the cells in 100 mL YPGA (Gal) medium (see above, but including 20 g/L galactose instead of D-glucose) and grown for another 15-18 h at 25°C and 160 rpm. The cells were harvested by centrifugation (7500 *g*, 10 min, 4°C), resuspended in 30 mL TEK buffer (50 mM Tris-HCl pH 7.5, 1 mM EDTA, 100 mM KCl) and centrifuged again. Then, the cells were carefully resuspended in 2 mL TES buffer (50 mM Tris-HCl pH 7.5, 1 mM EDTA, 600 mM sorbitol; freshly added: 10 g/L bovine serum fraction V protein and 1.5 mM β-mercaptoethanol) and disrupted by shaking five times for 1 min with glass beads (0.45–0.50 mm diameter, Sigma-Aldrich). The crude extracts were recovered by washing the glass beads four times with 5 ml TES. The combined washes were centrifuged (7500 *g*, 10 min, 4°C), and the supernatant containing the microsomes was transferred into an ultracentrifuge tube. After ultracentrifugation (100,000 *g*, 90 min, 4°C), the supernatant was carefully removed and the microsomal pellet was gently washed with 2.5 mL TES buffer, then with 2.5 mL TEG buffer (50 mM Tris-HCl pH 7.5, 1 mM EDTA, 30% glycerol). The microsomal fractions were homogenized in 2 mL TEG buffer using a glass homogenizer (Potter-Elvehjem, Carl Roth) and aliquots were stored at −20°C until further use.

### Enzyme assays

IDS assays were carried out using 3 µg of purified ApIDS protein desalted in IDS assay buffer (25 mM MOPSO, pH 7.2, 10% (v/v) glycerol), 1 mM metal ion cofactor (MgCl2, MnCl2, or CoCl2), and the substrates IPP and DMAPP (each 50 µM) in 100 µL assay buffer at 30 °C for 1 h. Product formation was monitored by liquid chromatography-tandem mass spectrometry (LC-MS/MS) as described below.

GES activity was determined in assays (total volume, 100 µL) containing 20 µL yeast microsomes harboring ApGES, 25 mM Tris-HCl (pH 7.5) and 50 µg/µl GPP. The assays were overlayed with 100 µL hexane and incubated for 20 min at 22°C. Enzyme products were extracted by vortexing for 1 min, and 1 µL of the hexane phase was injected into the GC-MS (see below).

For measuring G8H activity, 10 µL microsomes harboring either ApG8H alone or in combination with the P450 reductase ApRed were incubated in 25 mM sodium phosphate buffer (pH 7.0) with 25 mM substrate (geraniol, geranial, citronellol, or citronellal, respectively) and 1 mM NADPH in a total volume of 100 µL for 2 h at 30°C. Assays were then overlayed with 100 µL ethyl acetate and vortexed for 1 min. G8H products were analyzed by injecting 1 µL of the ethyl acetate phase into the GC-MS. The indole hydroxylase BX2 that is involved in benzoxazinoid formation in maize (26) was used as a negative control. Screening ApG8H activity with other substrates including citral A+B, nerol, linalool, limonene, and myrcene was performed by adding 10 µL of the substrate (0.5 mM dissolved in methanol) to 500 µL of living yeast cells induced with galactose-containing medium (see above). Cells were further incubated for 24 h at 28°C and 200 rpm, and afterwards extracted with 200 µl ethyl acetate. An aliquot (1 µL) of the organic phase was injected into GC-MS for enzyme product analysis.

HGO activity was analyzed using assays containing 40 µg purified protein, 1 mM NAD^+^ or NADP^+^, respectively, and 0.5 mM substrate (8-hydroxygeraniol, geraniol, nerol, or β-citronellol) in a total volume of 50 µL MOPS buffer (0.1 M). Assays were overlayed with 200 µL ethyl acetate, incubated for 2 h at 30°C, and enzyme products were extracted by vortexing the assay for 1 min. One µl of the organic phase was injected into GC-MS for enzyme product analysis.

ISY activity was measured in assays (total volume, 50 µL) containing 20 µL microsomes, 50 mM MOPS pH 7.5, 1 mM NADPH, and 0.5 mM 8-oxogeraniol. Assays were incubated for 2 h at 30°C, overlayed with 100 µL ethyl acetate, and products were extracted by vortexing the assays for 1 min. One µl of recombinant and purified CrISY from *Catharanthus roseus* (8) was tested under the same conditions as described above either in the presence or absence of 20 µL yeast microsomes as negative control.

NEPO activity was determined as described above for HGO with 3 µg of purified protein and 0.5 mM 7*S*-*cis-trans*-nepetalactol as substrate.

### Gas chromatography-mass spectrometry analysis

Qualitative analysis of volatile sex pheromones released from sexual female pea aphids was conducted using an Agilent 6890 Series gas chromatograph coupled to an Agilent 5973 quadrupole mass selective detector (Agilent Technologies, Santa Clara, CA, USA; injector temperature, 220°C; interface temp, 250°C; quadrupole temp, 150°C; source temp, 230°C; electron energy, 70 eV). The constituents of the volatile bouquet were separated using a ZB5 column (Phenomenex, Aschaffenburg, Germany; 30 m× 0.25 mm × 0.25 µm) and He as carrier gas (2 mL/min). The SPME sample was injected without split at an initial oven temperature of 70°C. The temperature was held for 2 min and then increased to 220°C with a gradient of 7°C min^−1^, and then further increased to 300°C with a gradient of 60°C min^−1^ and a hold of 2 min. Enzyme products extracted in hexane or ethyl acetate were analyzed using the same GC-MS system with a carrier gas flow of 1.5 mL min^−1^, splitless injection (1 µL sample) and a temperature program from 60°C (2 min hold) at 10°C min^-1^ to 220°C, and a further increase to 300°C with a gradient of 100°C min^-1^ and a hold of 2 min. Compounds were identified by comparison of retention times and mass spectra to those of authentic standards, or by comparison with reference spectra in the Wiley and National Institute of Standards and Technology libraries.

### Liquid chromatography-tandem mass spectrometry analysis of IDS products

ApIDS products GPP, FPP, and GGPP were analyzed as recently described in Lackus et al. (42) using an Agilent 1200 HPLC system (Agilent Technologies, Santa Clara, CA, USA) coupled to an API 6500 triple-quadrupole mass spectrometer (Applied Biosystems, Foster City, CA, USA). For separation, a ZORBAX Extended C-18 column (1.8 μm, 50 mm × 4.6 mm; Agilent Technologies, Santa Clara, CA, USA) was used. The mobile phase consisted of 5 mM ammonium bicarbonate in water as solvent A and acetonitrile as solvent B, with the flow rate set at 0.8 mL/min and the column temperature kept at 20°C. Separation was achieved by using a gradient starting at 0% B (v/v), increasing to 10% B in 2 min, 64% B in 12 min, and 100% B in 2 min (1-min hold), followed by a change to 0% B in 1 min (5-min hold) before the next injection. The injection volume for samples and standards was 1 μL. The mass spectrometer was used in the negative electrospray ionization (EI) mode. Multiple-reaction monitoring (MRM) was used to monitor analyte parent ion-to-product ion formation: *m/z* 312.9/79 for GPP, *m/z* 380.9/79 for FPP, and *m/z* 449/79 for GGPP.

### Phylogenetic analysis

Amino acid alignments were constructed using the MUSCLE algorithm (gap open, −2.9; gap extend, 0; hydrophobicity multiplier, 1.2; clustering method, UPGMB) implemented in MEGA7 (43). Tree reconstruction was done with MEGA7 using a Maximum Likelihood algorithm (model/method, given in the respective figure legends; substitutions type, amino acids; rates among sites, uniform rates; gaps/missing data treatment, partial deletion; site coverage cutoff, 80%). Bootstrap resampling analyses with 1000 replicates were performed to evaluate the tree’s topologies.

### Statistical analysis

Differences in phosphatase activities between the empty vector control and ApGES were compared with the Welch two sample t-test. Data were analyzed with R 4.2.0 (https://www.R-project.org/).

## Acknowledgments

We thank Michael Reichelt and Maritta Kunert for help with LC-MS/MS and GC-MS analysis, respectively, Nestor Jose Hernandez-Lozada for providing purified recombinant CrISY protein, Kimberly Falk for drawing the aphid cartoons shown in Figure 1, Julie Jaquiery for helpful advice regarding the generation of sexual femal aphids, and Elke Goschala, Claudia Pakebusch and Andreas Weber from the greenhouse team of the MPI-CE for rearing *Vicia faba* plants. We gratefully acknowledge Fen Li for preliminary experiments. This work was supported by the Max Planck Society. Additional support was provided by the ERC (778301).

## Notes

### Competing Interest Statement

The authors have declared no competing interest.

