## Supporting Information for "Biosynthesis of iridoid sex pheromones in aphids"

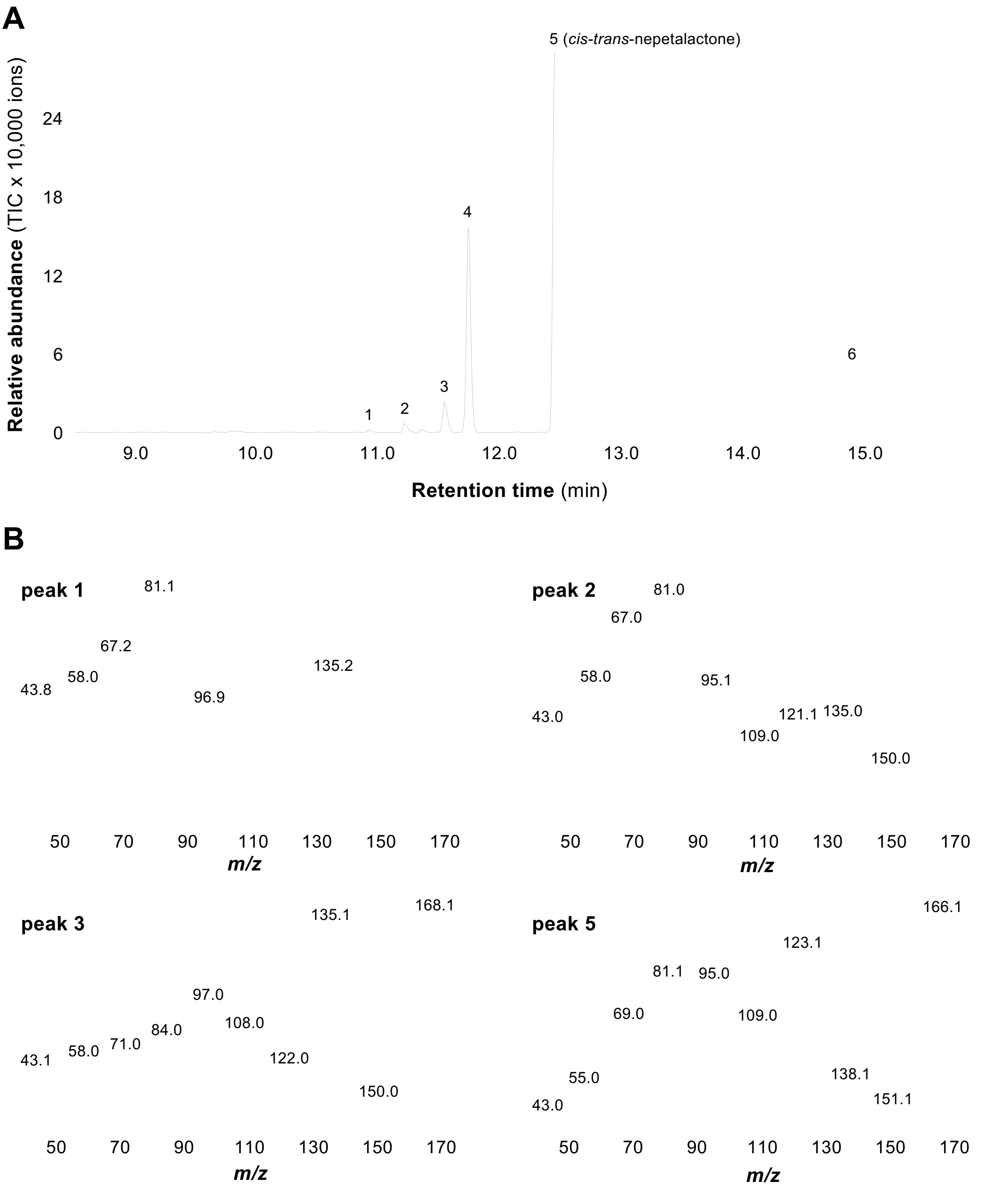

**Supplemental Figure S1: Sexual females of the pea aphid *Acyrthosiphon pisum* release volatile iridoids. (A)** Asexual *A. pisum* aphids were reared on *Vicia faba* plants in closed cellophane bags under short-day conditions and reduced temperature to stimulate the development of sexual females. Volatiles released from sexual female aphids were collected from the headspace of the plant/aphids using solid-phase microextraction (SPME) and analyzed with gas chromatography-mass spectrometry (GC-MS). 1, unidentified; 2, unidentified; 3, *cis-trans*-nepetalactol; 4, dodecamethylcyclohexasiloxane (contamination); 5, *cis-trans*-nepetalactone; 6, tetradecamethylcycloheptasiloxane (contamination). **(B)** Mass spectra of peaks 1,2,3, and 5 of the chromatogram shown in panel A.

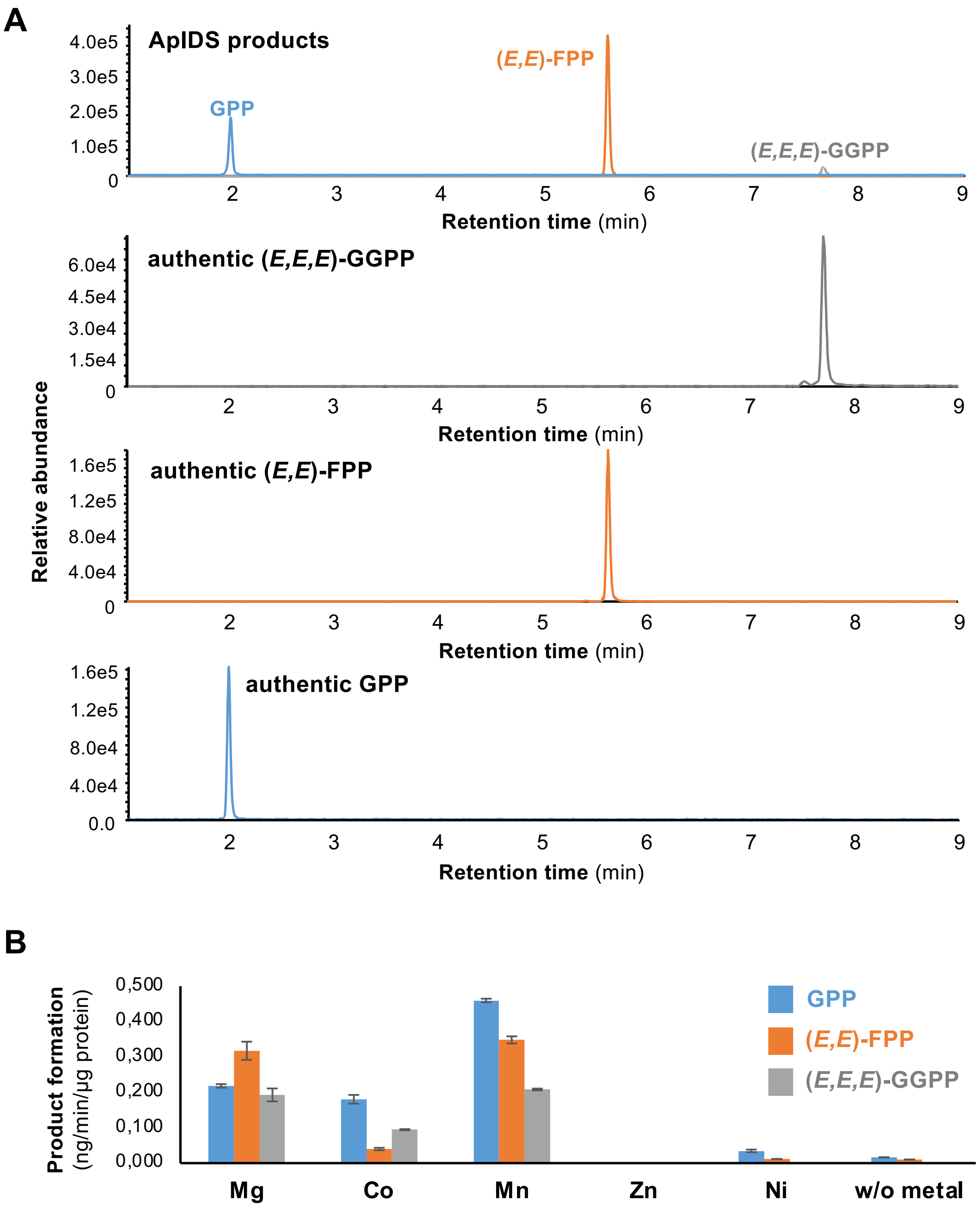

**Supplemental Figure S2: The metal ion cofactor influences product specificity of ApIDS. (A)** Identification of ApIDS products using authentic standard compounds. N-terminal truncated ApIDS was expressed as N-terminal His-tag-fusion protein in *Escherichia coli*, purified, and incubated with the substrates IPP and DMAPP in the presence of 1 mM $MgCl_2$. Products were analyzed using liquid chromatography-tandem mass spectrometry (LC-MS/MS) and identified with authentic GPP, (*E,E*)-FPP, and (*E,E,E*)-GGPP. **(B)** For testing the influence of the metal ion cofactor on product specificity, purified ApIDS was incubated with IPP, DMAPP, and 1 mM of cofactor, and product formation was analyzed using LC-MS/MS. Quantification of enzyme products was performed using standard curves made from authentic GPP, (*E,E*)-FPP, and (*E,E,E*)-GGPP. Means and SE are shown (n = 3 technical replicates).

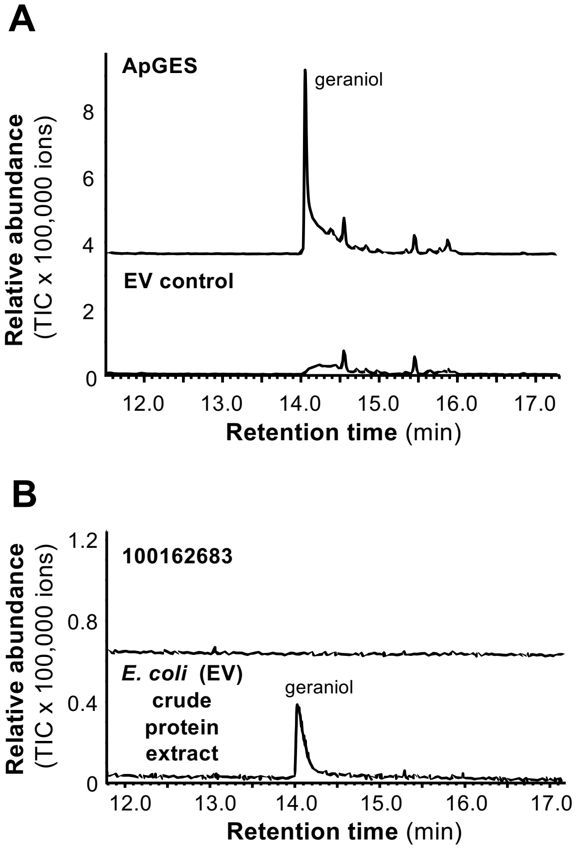

**Supplemental Figure S3: ApGES showed phosphatase activity with GPP. (A)** The phosphatase ApGES was expressed in *Saccharomyces cerevisiae* and microsomes harboring the recombinant protein were incubated with GPP. Geraniol was extracted with hexane and analyzed using gas chromatography-mass spectrometry (GC-MS). Microsomes prepared from *S. cerevisiae* carrying the empty expression vector were used as negative control. **(B)** Another potential phosphatase candidate (gene ID 100162683) annotated as inositol polyphosphate 1-phosphatase was expressed in *Escherichia coli,* purified, and incubated with GPP. A crude protein extract made from *E. coli* carrying the empty expression vector and fed with GPP showed unspecific phosphatase activity and was used as positive control. Reaction products were extracted with hexane and analyzed using GC-MS.

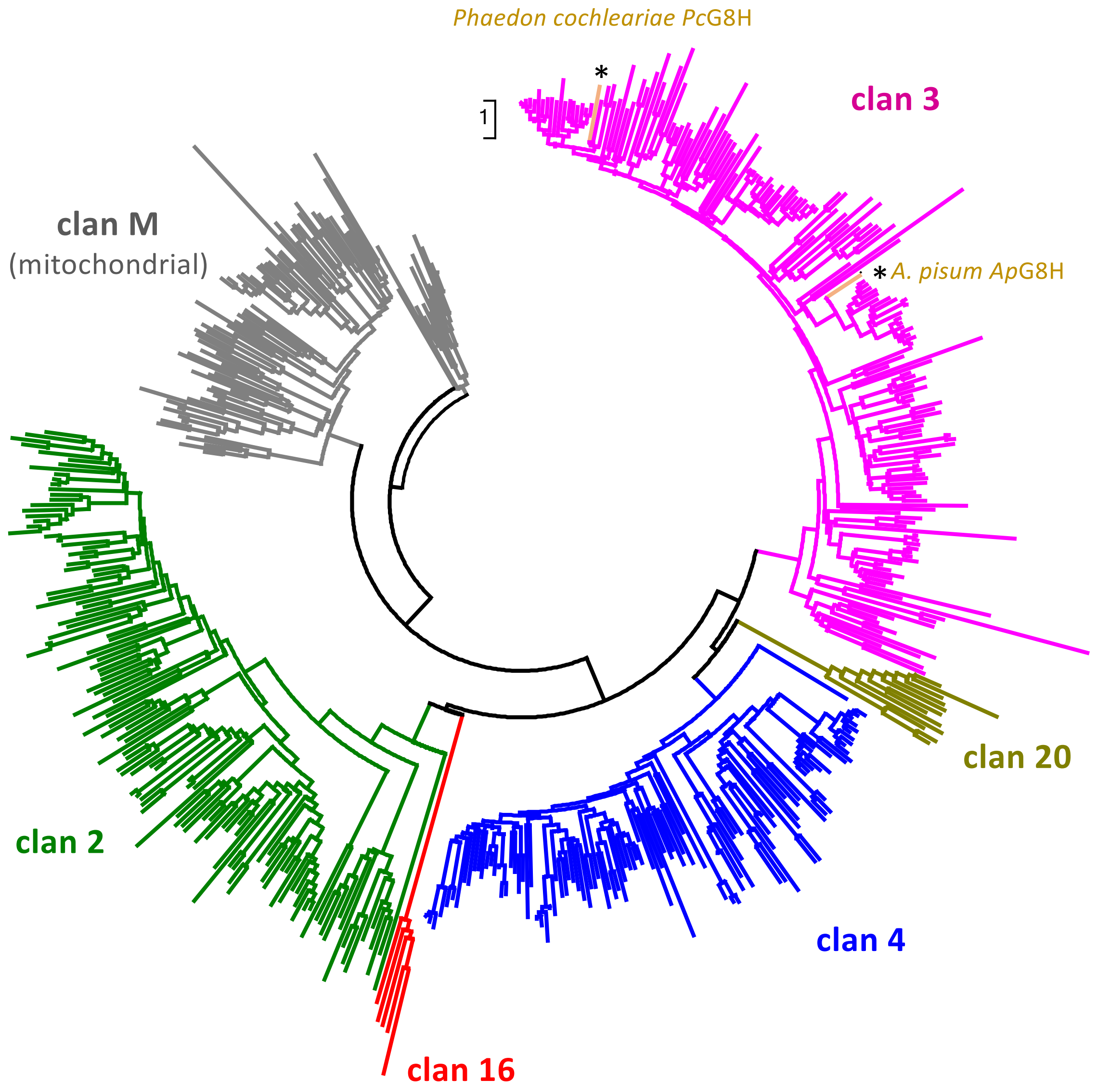

**Supplemental Figure S4: ApG8H from *A. pisum* and PcG8H from *P. cochleariae* both belong to the clan 3 of Arthropode P450s, but share only 35% amino acid similarity.** Cladogram analysis of Arthropode P450 proteins including ApG8H from *A. pisum* and PcG8H from *P. cochleariae*. The tree was inferred by using the Neighbor-Joining method based on the JTT matrix-based method. The rate variation among sites was modeled with a gamma distribution (shape parameter = 1). The analysis involved 542 amino acid sequences. All positions with less than 80% site coverage were eliminated. The tree is drawn to scale, with branch length measured in the number of amino acid substitutions per site.

**A**

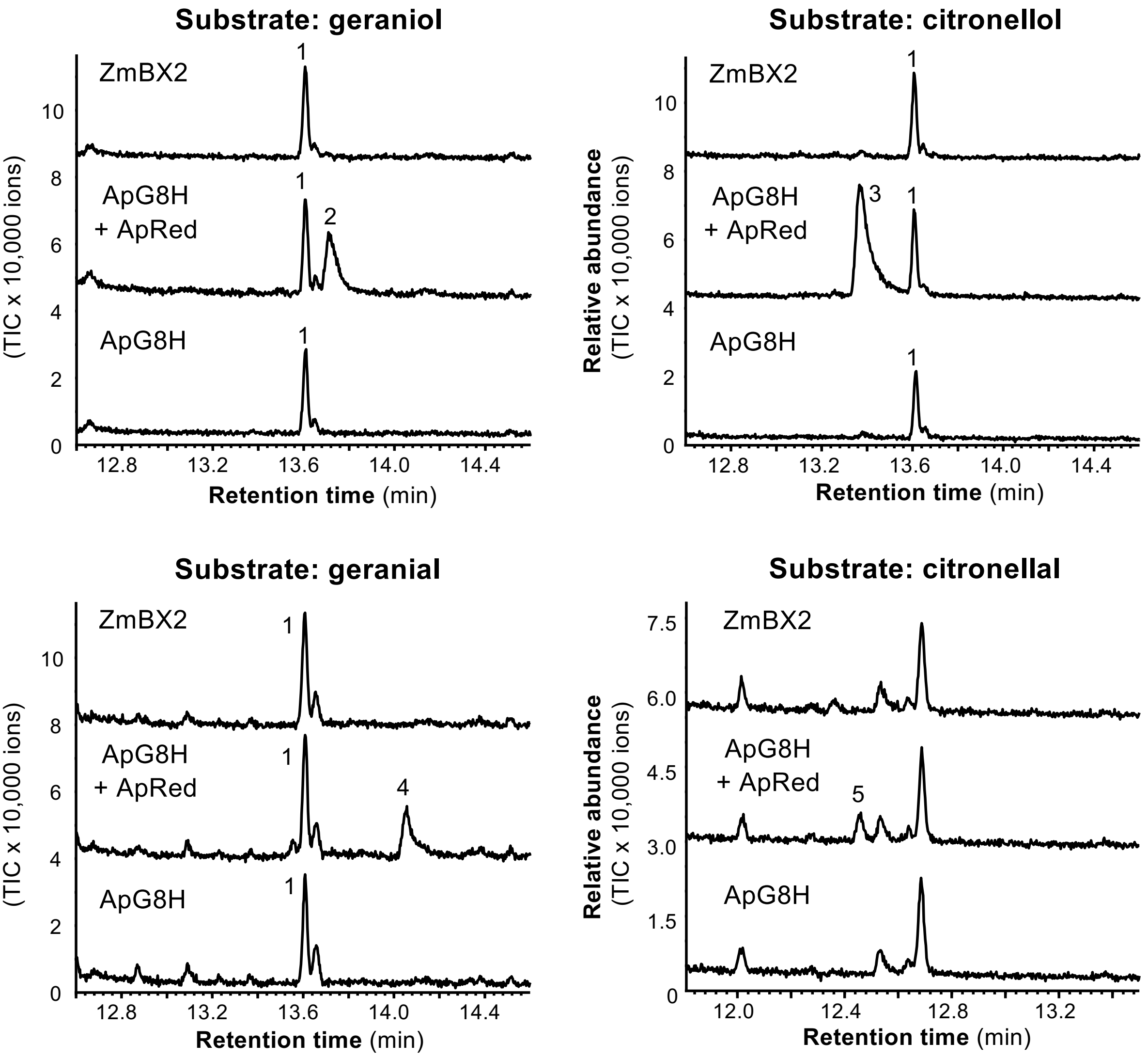

**B**

| | ApG8H | ApG8H + ApRed |
|---|---|---|
| geraniol | - | + |
| citral A+B | - | + |
| nerol | - | + |
| citronellol | - | + |
| linalool | - | + |
| limonene | - | - |
| myrcene | - | - |

**Supplemental Figure S5: Biochemical characterization of ApG8H. (A)** Yeast (*Saccharomyces cerevisiae*) microsomes containing either ApG8H, ApG8H in combination with the P450 reductase ApRed, or maize ZmBX2 as negative control were assayed with potential terpenoid substrates and NADPH as cosubstrate. Reaction products were analyzed using GC-MS. 1, di-tert-butylphenol (contamination); 2, 8-hydroxygeraniol; 3, 8-hydroxycitronellol; 4, 8-hydroxygeranial; 5, 8-hydroxycitronellal. Compounds were identified by comparisons with authentic standards (8-hydroxygeraniol) or the NIST, WILEY, and Adams mass spec libraries. **(B)** *S. cerevisiae* liquid cultures expressing either *ApG8H* alone or in combination with *ApRed* were fed with potential terpenoid substrates. Reaction products were extracted with ethylacetate from the cultures and analyzed using GC-MS. +, activity could be observed; -, no activity.

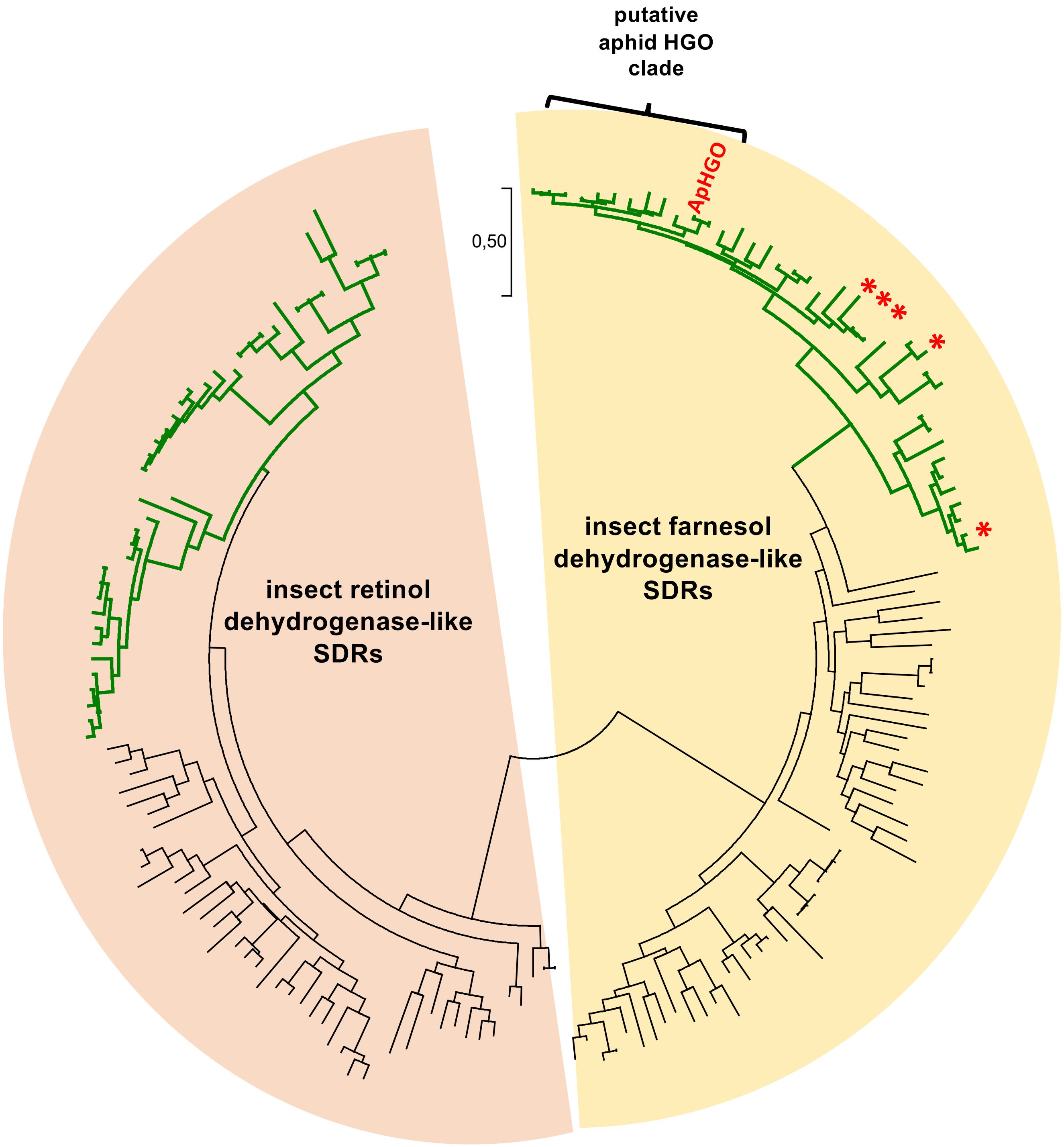

**Supplemental figure S6: ApHGO belongs to the aphid clade of farnesol dehydrogenase-like short-chain dehydrogenases/reductases (SDRs).** ApHGO is shown in red and other farnesol dehydrogenase-like SDRs from *A. pisum* are marked with red asterisks. The tree was inferred by using the Maximum Likelihood method based on the JTT matrix-based model and is drawn to scale, with branch lengths measured in the number of amino acid substitutions per site. Aphid proteins are marked with green branches.

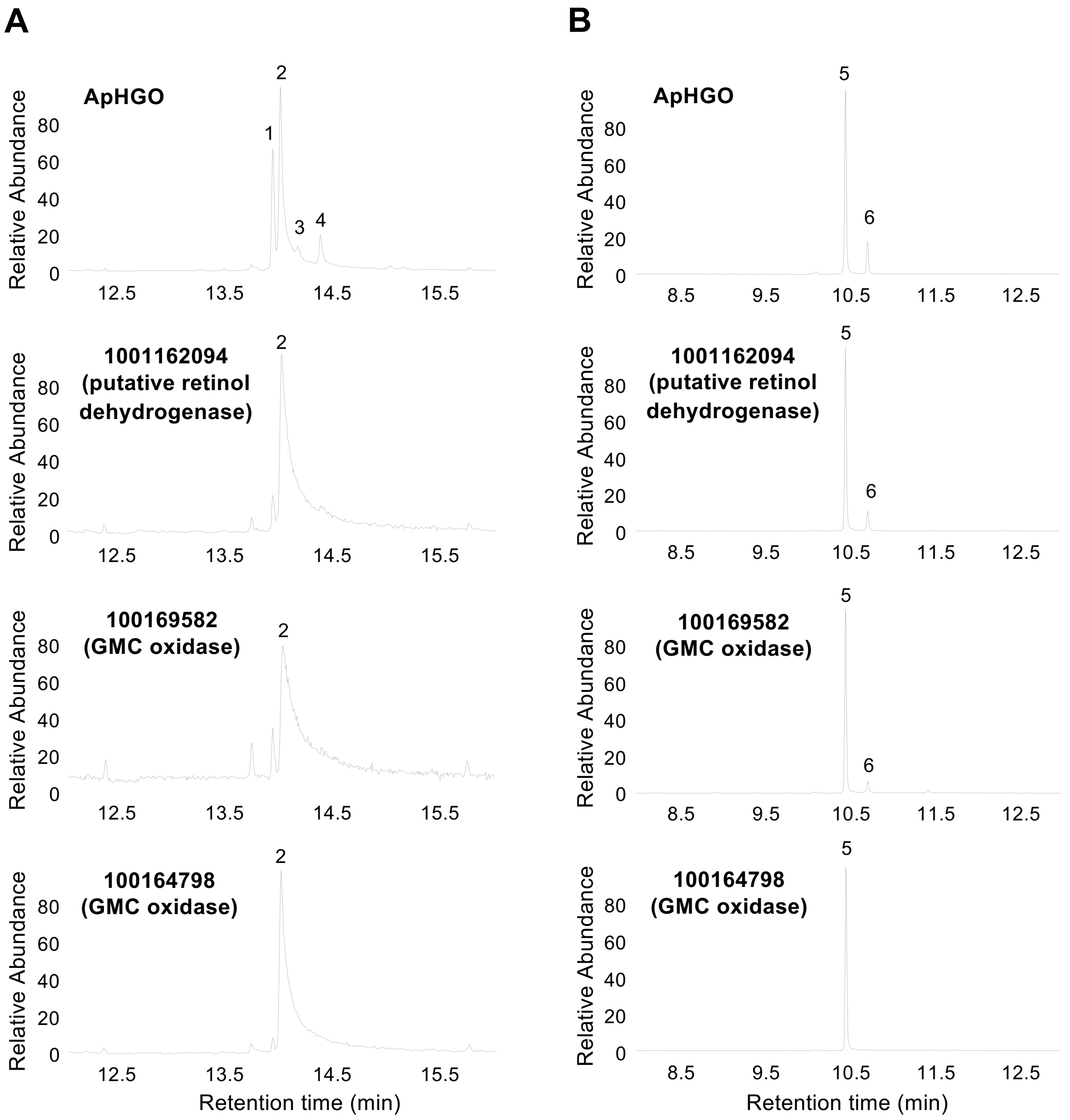

**Supplemental Figure S7: The putative retinol dehydrogenase 1001162094 and the GMC oxidase 100169582 (ApNEPO) accepted geraniol but not 8-hydroxygeraniol as substrate.** Enzymes were expressed as N-terminal His-tag-fusion proteins in *Escherichia coli*, purified, and incubated with 8-hydroxygeraniol **(A)** or geraniol **(B)** in the presence of NADP. Enzyme products were extracted from the assays and analyzed using gas chromatography-mass spectrometry. 1, 8-oxogeraniol; 2, 8-hydroxygeraniol; 3, 8-oxogeranial; 4, 8-hydroxygeranial; 5, geraniol; 6, geranial. The GMC oxidase 100164798 showed no activity with the tested substrates.

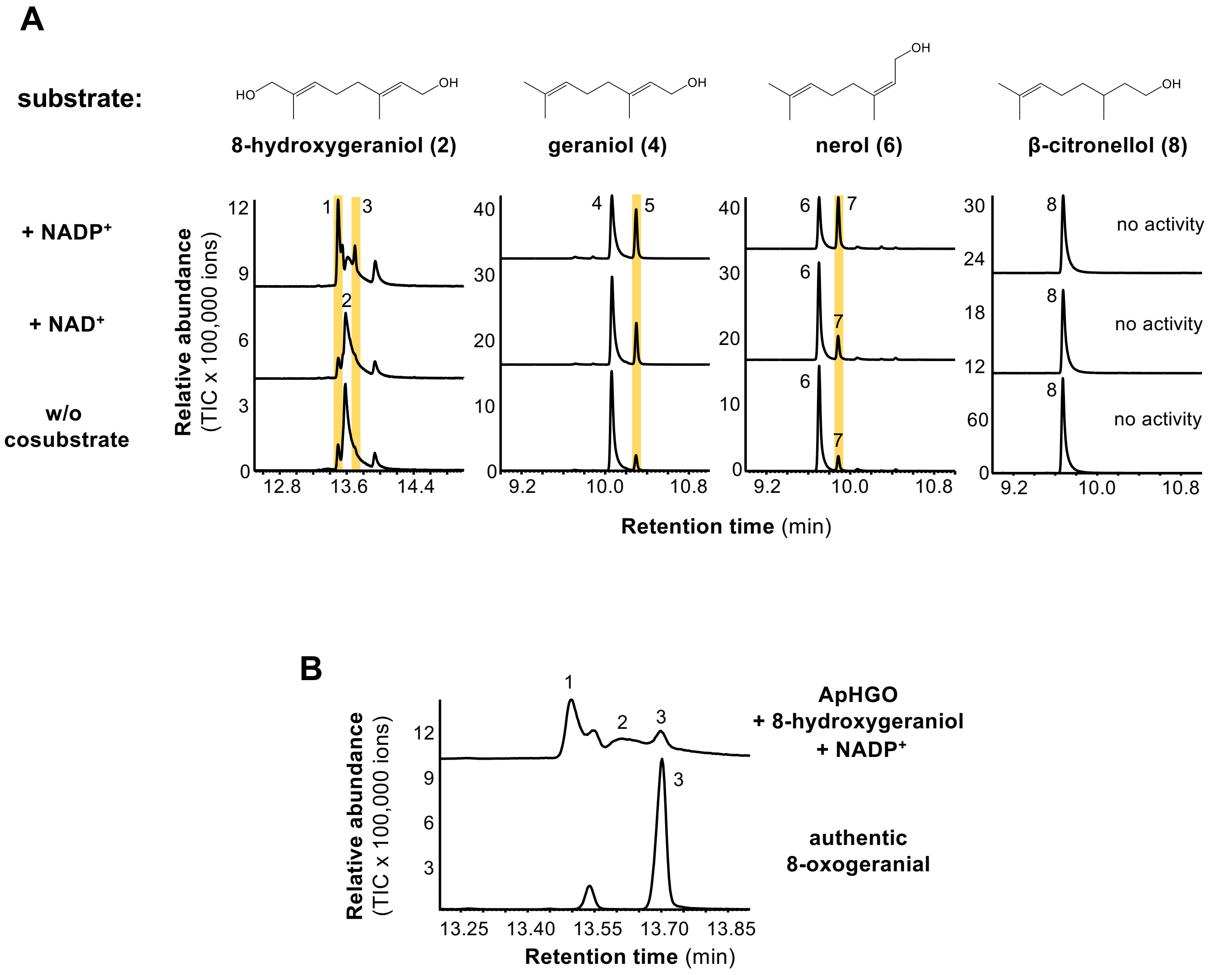

**Supplemental Figure S8: Biochemical characterization of ApHGO. (A)** ApHGO was expressed as N-terminal His-tag-fusion protein in *Escherichia coli*, purified, and incubated with potential terpenoid substrates either in the absence or presence of NAD(P). Enzyme products were extracted from the assays and analyzed using gas chromatography-mass spectrometry. 1, 8-oxogeraniol (partially oxidized product); 2, 8-hydroxygeraniol; 3, 8-oxogeranial (fully oxidized product); 4, geraniol; 5, geranial; 6, nerol; 7, neral; 8, β-citronellol. **(B)** The ApHGO reaction product 8-oxogeranial were identified using an authentic standard.

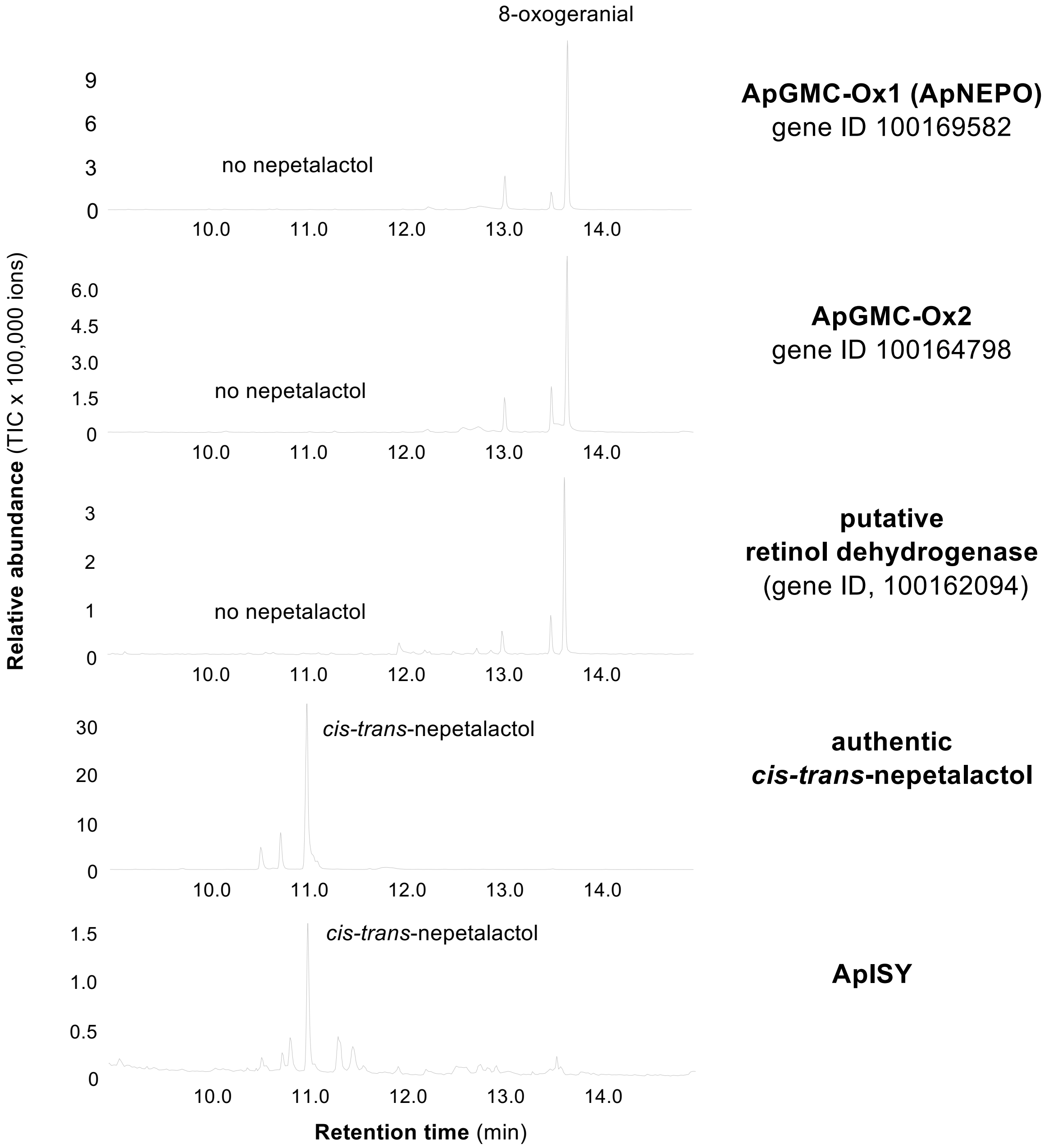

**Supplemental Figure S9: Characterization of *A. pisum* oxidoreductase candidate enzymes.** The two GMC oxidases and the putative retinol dehydrogenase were expressed as N-terminal His-tag-fusion proteins in *Escherichia coli*, purified, and incubated with 8-oxogeranial and NADPH. ApISY was expressed in *Saccharomyces cerevisiae* and yeast microsomes harboring the recombinant protein were fed with 8-oxogeranial and NADPH. Reaction products were extracted from the assays and analyzed using gas chromatography-mass spectrometry.

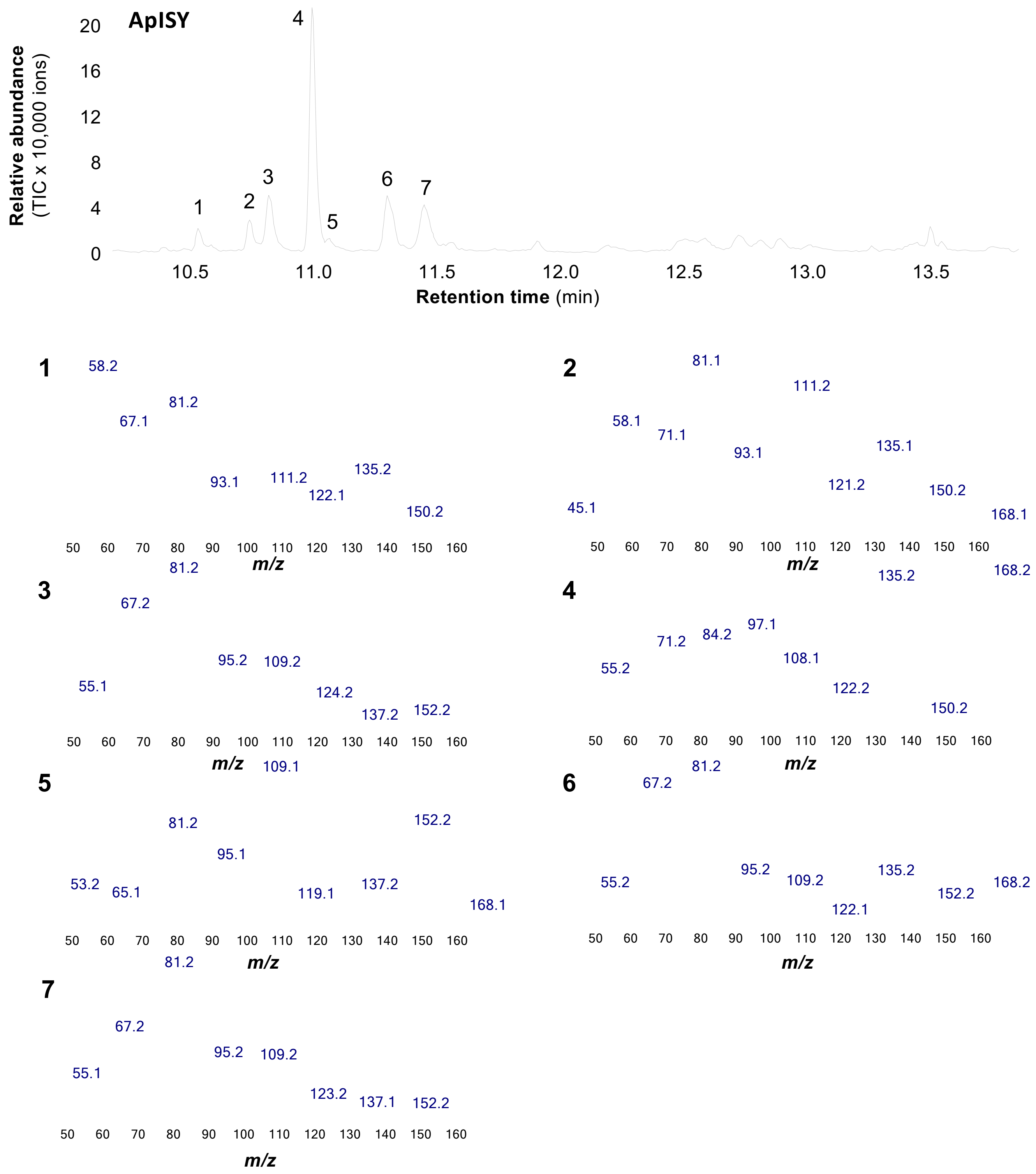

**Supplemental Figure S10: Mass spectra of the ApISY reaction products.** Yeast (*Saccharomyces cerevisiae*) microsomes containing ApISY were assayed with 8-oxogeranial as substrate and NADPH as cosubstrate. Reaction products were extracted with ethylacetate and analyzed using gas chromatography-mass spectrometry. 1, *cis-trans*-iridodial; 2, unidentified; 3, unidentified; 4, *cis-trans*-nepetalactol; 5, unidentified; 6, unidentified; 7, unidentified.

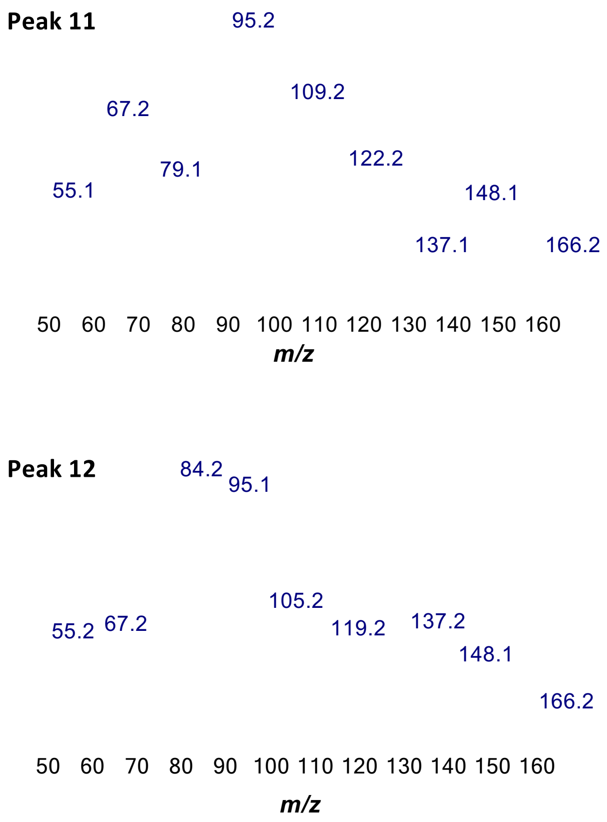

**Supplemental figure S11: Mass spectra of peaks 11 and 12 in Figure 4.**

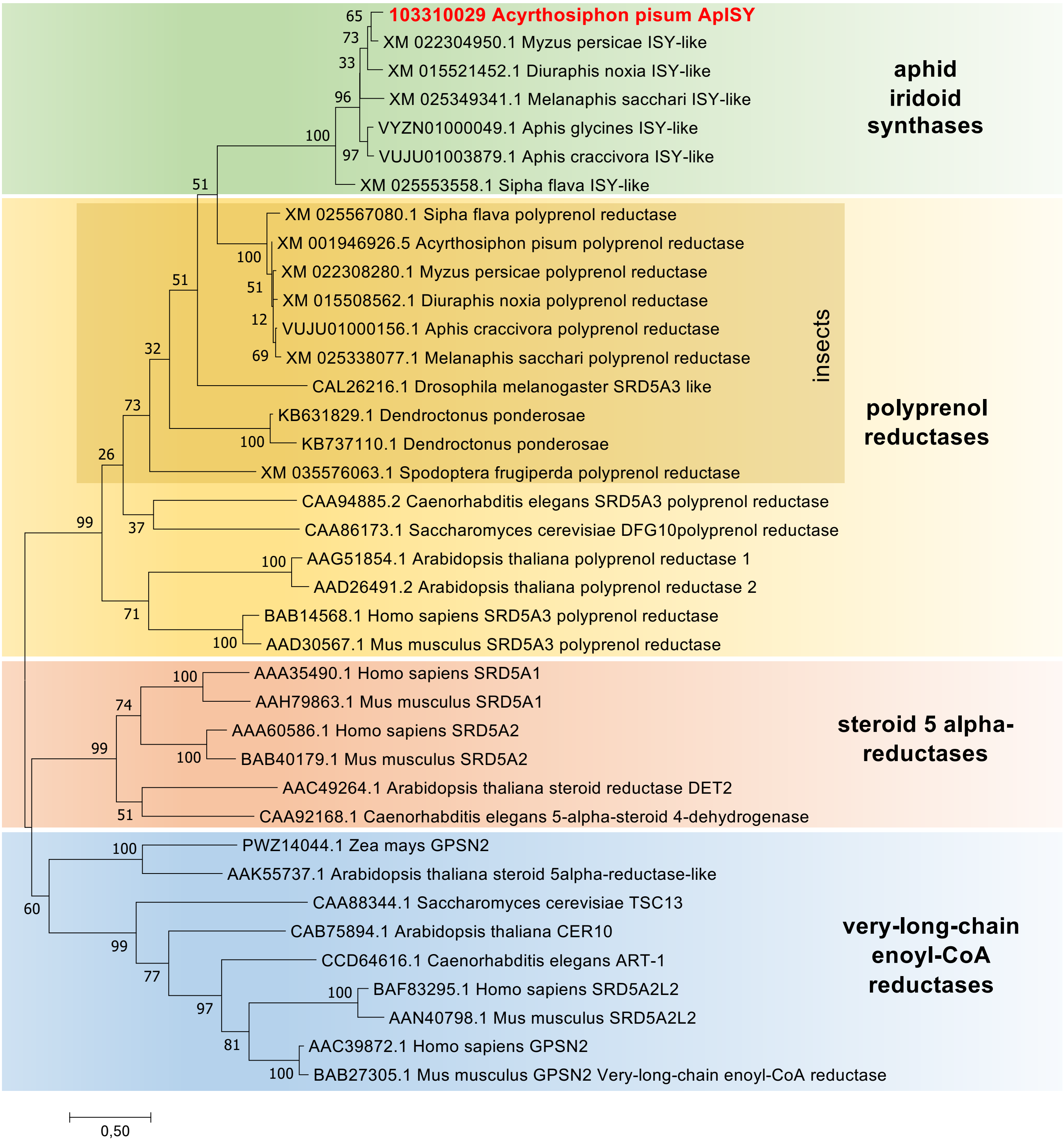

**Supplemental Figure S12: ApISY belongs to the steroid-5-alpha reductase-like family (SRD5A-like).** The tree was inferred by using the Maximum Likelihood method based on the JTT matrix-based model and is drawn to scale, with branch lengths measured in the number of amino acid substitutions per site. All positions with less than 80% site coverage were eliminated. Bootstrap values are given next to each node (n = 1000).

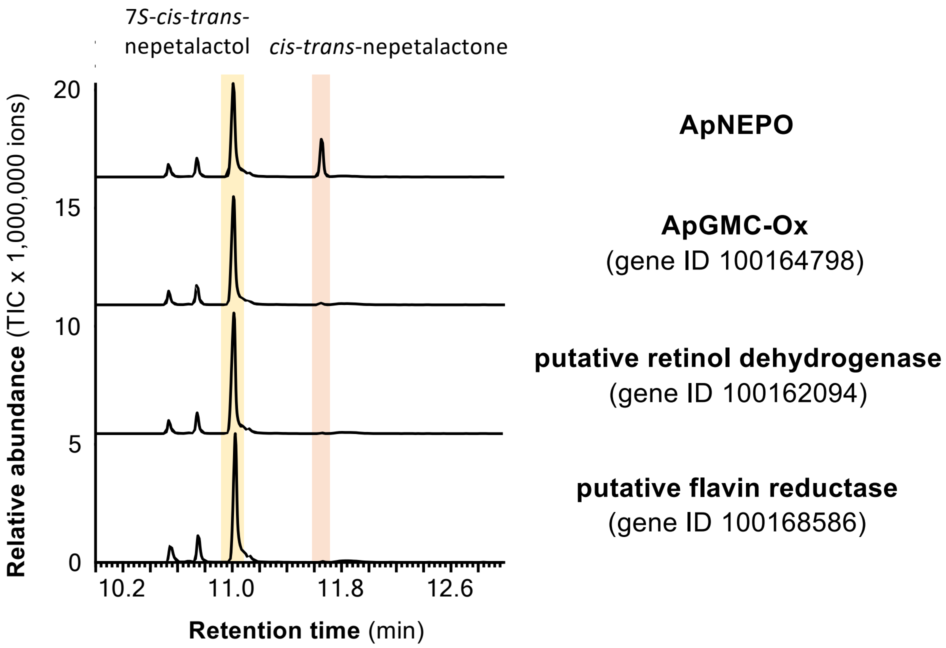

**Supplemental Figure S13: Biochemical characterization of ApNEPO.** ApNEPO and three other putative oxidoreductases specifically expressed in hind legs of female *A. pisum* were expressed as N-terminal His-tag-fusion proteins in *Escherichia coli*, purified, and incubated with 7*S*-*cis*-*trans*-nepetalactol in the presence of NADP. Enzyme products were extracted from the assays with ethylacetate and analyzed using gas chromatography-mass spectrometry.

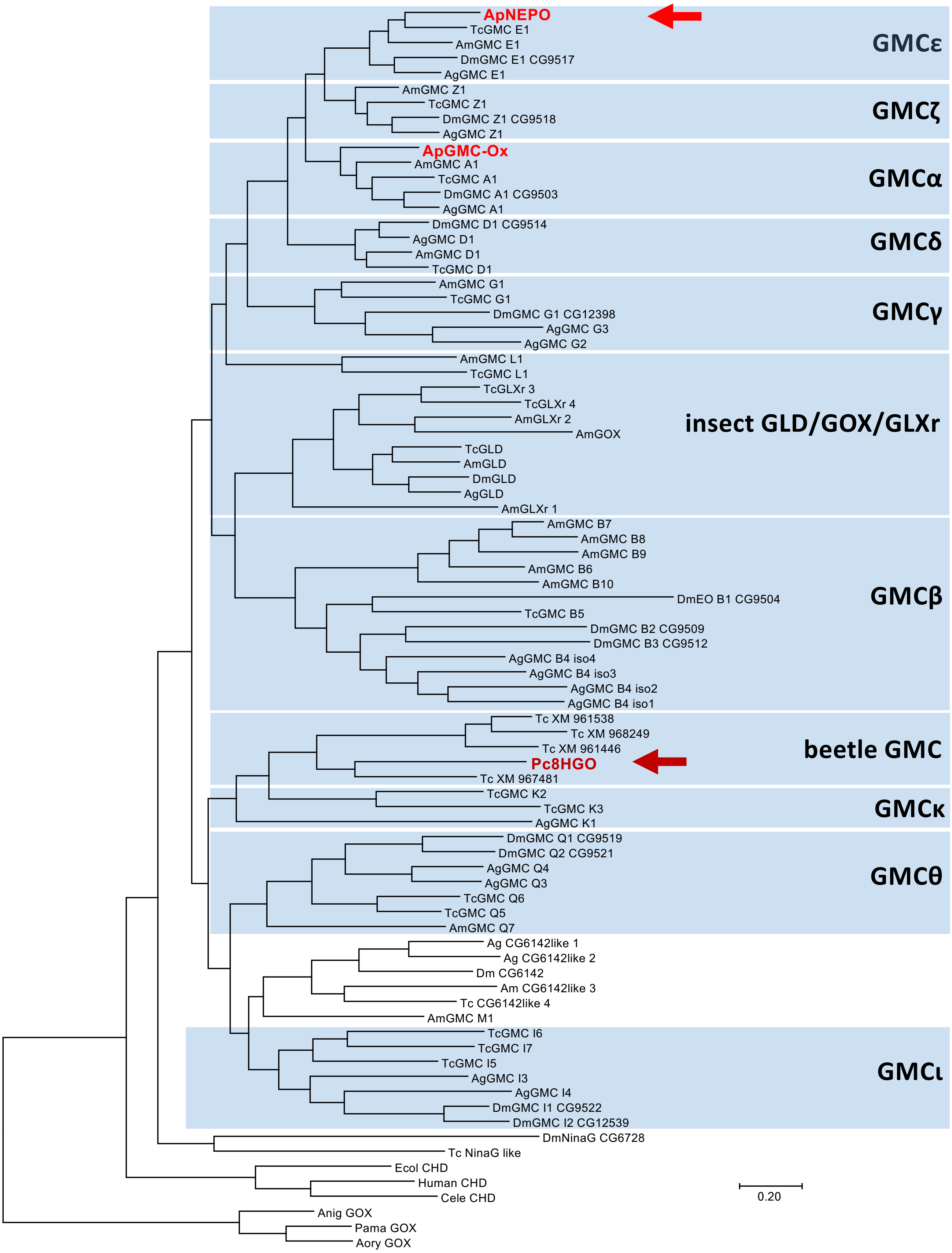

**Supplemental figure S14: ApNEPO belongs to the GMCε clade of GMC oxidases and is not related to Pc8HGO from *Phaedon cochleariae*.** The tree was inferred by using the Maximum Likelihood method based on the JTT matrix-based model and is drawn to scale, with branch lengths measured in the number of amino acid substitutions per site.

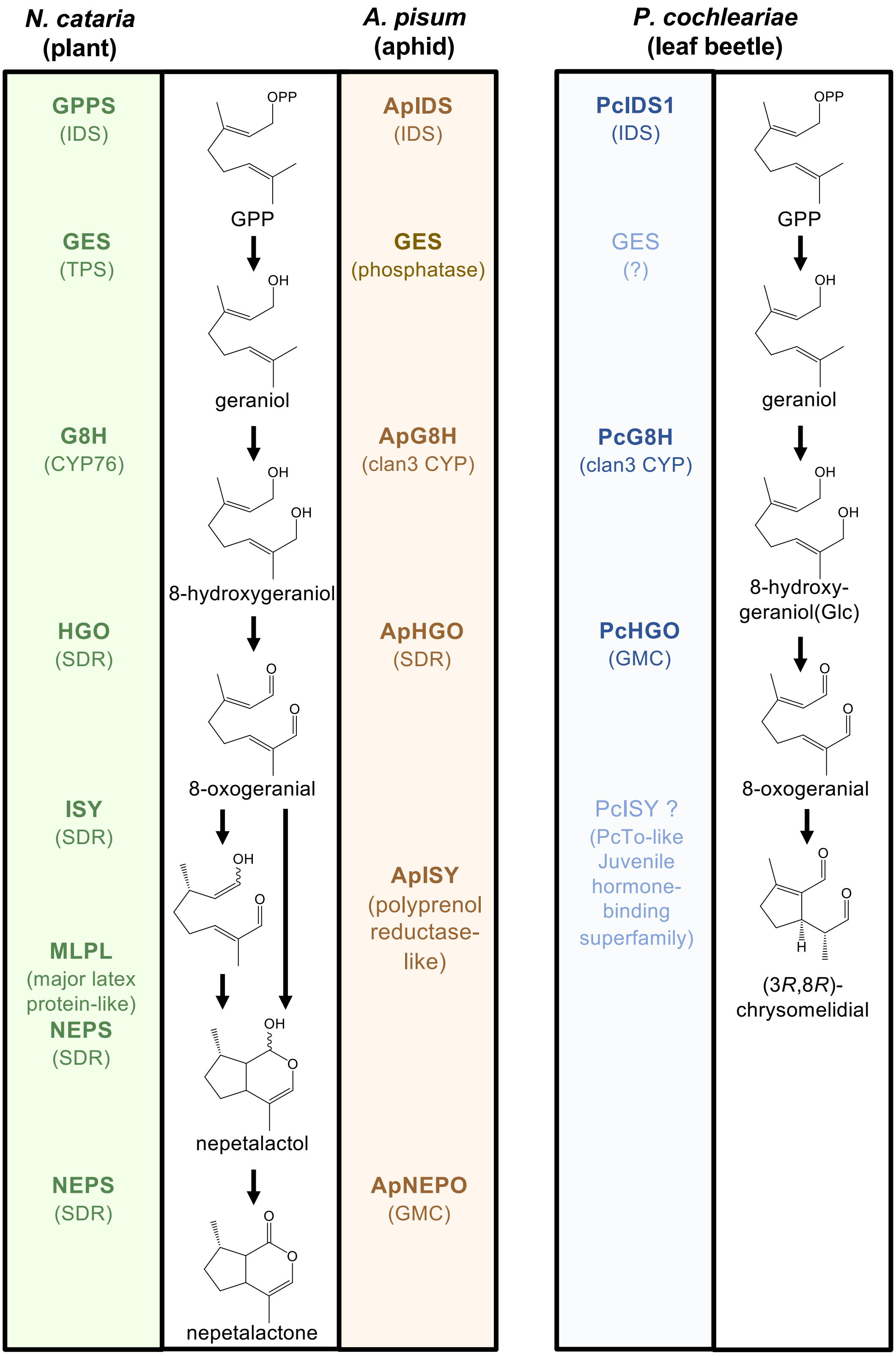

**Supplemental Figure S15: The iridoid pathways in plants, aphids, and beetles evolved independently from each other.** While the plant and aphid pathways both lead to nepetalactol, the beetle *Phaedon cochleariae* produces chrysomelidial. Enzymes identified and characterized are shown in bold. IDS, isoprenyldiphosphate synthase; TPS, terpene synthase; CYP, cytochrome P450 monooxygenase; SDR, short-chain dehydrogenase/reductase; GMC, glucose-methanol-cholin oxidoreductase.

**Table S1: Genes selectively expressed in hind legs of sexual female pea aphids.**

The table is attached as Excel file

**Table S2: Expression of mevalonate and nepetalactone pathway genes in hind legs and front legs of different sexual stages of *A. pisum*.** RNA was extracted from aphid legs and sequenced, and the obtained reads were mapped onto the *A. pisum* genome version v3. Mean RPKM values are shown (n = 3). f-hl, hind legs of sexual females; f-fl, front legs of sexual females; af-hl, hind legs of asexual females; m-hl, hind legs of males.

| Gene ID | Original gene annotation | f-hl | f-fl | af-hl | m-hl |
|---|---|---|---|---|---|
| 100162815 | acetyl-CoA acetyltransferase | 1298.52 | 9.64 | 11.61 | 8.27 |
| 100165154 | HMG-CoA synthase | 5143.01 | 3.72 | 4.46 | 5.07 |
| 100165462 | HMG-CoA reductase | 1502.11 | 18.64 | 16.87 | 17.87 |
| 100163305 | mevalonate kinase | 374.42 | 37.29 | 28.55 | 25.64 |
| 100574505 | phosphomevalonate kinase | 1447.09 | 21.88 | 26.33 | 39.36 |
| 100163413 | phosphomevalonate kinase-like | 230.95 | 4.26 | 4.66 | 6.32 |
| 100158798 | diphosphomevalonate decarboxylase | 2093.95 | 29.54 | 36.26 | 31.61 |
| 100166744 | isopentenyl diphosphate isomerase | 2136.28 | 44.48 | 29.17 | 34.51 |
| 100144905 | isoprenyl diphosphate synthase (ApIDS) | 395.83 | 20.26 | 13.31 | 13.90 |
| 100158803 | dolichyldiphosphatase 1 (ApPhos) | 122.32 | 5.92 | 10.20 | 8.48 |
| 100162683 | inositol polyphosphate 1-phosphatase | 19.23 | 0.91 | 0.78 | 0.97 |
| 100165972 | P450 (ApG8H) | 3409.27 | 13.19 | 24.46 | 48.41 |
| 100160284 | P450 reductase (ApG8H reductase) | 804.93 | 91.65 | 159.00 | 162.16 |
| 100301633 | farnesol dehydrogenase (ApHGO) | 10416.49 | 219.39 | 1062.13 | 1207.17 |
| 100162094 | retinol dehydrogenase | 96.65 | 12.37 | 0.52 | 0.48 |
| 103310029 | polyprenol reductase (ApISY) | 140.59 | 0.00 | 0.03 | 0.05 |
| 100169582 | GMC oxidase (ApNEPO) | 28.53 | 0.00 | 0.00 | 0.03 |
| 100164798 | GMC oxidase | 3382.68 | 0.03 | 0.14 | 0.67 |
| 100168586 | flavin reductase | 266.86 | 31.77 | 33.21 | 32.24 |

**Table S3: Signal peptide prediction with TargetP.** Prediction probabilities signal peptides and the predicted cleavage sites are shown. mTP, mitochondrial transfer peptide; SP, signal peptide; CS, cleavage site.

| Protein | Prediction | SP | mTP | CS Position |
|---|---|---|---|---|
| ApIDS | mTP | 0.001316 | **0.805817** | CS pos: 33-34 |
| ApGES | - | 0.000025 | 0.000236 | |
| ApG8H | - | 0.265424 | 0.010640 | |
| ApRed | - | 0.000022 | 0.000003 | |
| ApHGO | - | 0.002556 | 0.000248 | |
| ApISY | - | 0.113837 | 0.004443 | |
| ApNEPO | - | 0.167099 | 0.082896 | |
| 100164798 | - | 0.041727 | 0.000675 | |
| 100162683 | mTP | 0.001071 | 0.507994 | CS pos: 23-24 |
| 100168586 | SP | 0.517806 | 0.004666 | CS pos: 20-21 |
| 100162094 | - | 0.000450 | 0.002174 | |

**Table S4: Sequences synthesized or amplified in this study.**

| Gene | Note | Sequence |
|---|---|---|
| ApIDS | N-terminal truncated<br><br>optimized for *E. coli* | ATGTCAACTGTTCGTGCCCCACCCGTCCCGCCCGTGATTACCGGTACCGCTGTCTCAAAGGACGAGACACG<br>TGACTTTATGGCGGTTTTCCCTGATGTAGTACGCGACCTCACCGATACCGGACGTAATCTGGACGTGCCGG<br>ACGTTACCAAATGGCTCGCCAAACTGTTACAATATAACGTGCCTGGCGGGAAGAAGAACCGCGGCCTGGCG<br>CTGGTCTTAAGCTATAAGATGTTAAGCTCGCCGGCAGATCAAACAGATGAGAATATCCGTCTGAGTTATAT<br>CTTGGGGTGGTGTGTGGAGATTCTGCAAGCGTATCAATTAGTTTTAGACGATATCATGGATAATGCAATCA<br>CGCGCCGTGGGCGTCCGTGCTGGTACCGCCATAATGATATTGGCCTGATGGCCGTTAATGATGGTGTCCTG<br>CTCGAACAGAGCATCTATCAACTCATCAAGAAGTACTTTAAGGACAAGCCTTATTATACTCATATCTTGGA<br>GCTTTTCTATGACGTTACCATGAAAACCTCTATGGGGCAATGTCTGGACATGCTTACCGCAAATTCTTTTA<br>AGAGTAAGAAGCTGGAGAAGTATACTATGGAGAATTACACAGCAATCGTCAAGTATAAGACGGCTTACTAT<br>TCGTTCTTCTTACCTGTGTGTTTAGCAATGCGCATGACTAATATTAATGACCCGGAGATCTTCCGTCAGGC<br>AAAGACCATCTTATTAGAGATGGGGCACTTCTTCCAAGTTCAAGATGACTTCCTGGACTGCTATGGCGATC<br>CGGACGTGATGGGTAAGATTGGAACCGACATTGAAGATGGCAAGTGTTCTTGGCTTGCGGTAGTCGCCCTT<br>CAGAAGGTTAATAGCGAGCAAAAGAAGATTATGGAAGACAATTATGGCATCGATAACCCCGCTAATGTAGC<br>AGTTATCAAGGACTTGTACGCGCAGTTGAAGTTACCCGACACTTTCCACTTGTACGAAGAGGAGAGCTATA<br>AGCTGATTTGTACTCACATCCAACAACTTAGCCGCGGCTTATCGCAGGATATGTTCTTCAAATTCCTGGAA<br>AAGATCTACAAACGCACCCTCTAA |
| ApGES | optimized for *S. cerevisiae* | ATGATGACTTACTCCGTTTCTTCTAATAAGAAGCATGACTTGTCTAACCGTAACATCTTTGGCAAGCTATT<br>AGCTCTTTTCTCATTAACTCCCTTTGTGATACTTTCTGGGTTCATCTCTTTGATATTATTTAGACGTGATT<br>TACACACCATCACGTTCTTCTTCGGTGTACTGTTAAATGAGATTTGTAACACGGTATTGAAGCACATTTTG<br>AGGGAACCCAGGCCTTTGGCAAGGAACACGAACCTATTATATAGCGAATACGGCATGCCAAGCTCTCATTC<br>CCAGTTTATGTGGTTCTTCGCCTCTTACATGCTATACTTTACGTTCATCAGGTTGCAGTACGCTAACAACA<br>AAGCATTCAAGGAGTTCTTTTGGAAGGTCGCTGGCGCAGTCAGCTGCATAGCTATAGCATGCATCGTTTCG<br>TACAGTCGTATTTTCTTGCAGTACCATACATGGAAGCAGGTTATATACGGTGCATTGTTTGGTATAATTAT<br>TGGAACCATTTGGTTCACAATTATCAATGTCGTATTGACTCCGTACTTCCCTACAGTTATCTCCTGGAAAA<br>TATCAGAGCTTTTCTTACTTAGGGACACTACTTTGATTCCTAACGTCCTTTGGTTCGAATATACAAATATC<br>AGACACGAGGCCGGAGCACGTGCCAGAAGAAGAAAGAGTATTTCCGCCAAGTCACAATAA |
| ApG8H | amplified from cDNA | ATGTTTGAATTCGTCTACGAACTGTTCGATCTGAAAATGCTTTTGGTCACCGCTTTTCTGGGTGCCATATA<br>CGTGTATTCCACATGGACCCACAGCCATTGGTCCAAGCTGGGCATATCCAGTCCGTCTGCCCCGGTGCCGT<br>TGTTCGGGCACGCGATGCCCTCCATGTTGGGACAGATGCACTTCATGGATGTGTTGCACAACCTTTACAAG<br>GAGCTGGGCGACCAAAGGTTTGGTGGCATTTACACAATGCGAACACCGCAGCTCCTCGTCAAAGACCCAGA<br>ACTAATAGGACACATACTGATCAAAGACTTCAACAATTTCACGGACCGCGGATTATACGCTGGCACACACA<br>CAAACCCGCTCAACAATAATATATTCTTCACACGAGGCGAACGATGGAAAACGATGCGGCAAAAGCTCAGT<br>CCCACATTCACGGCCAACAAACTGAAGTACATGAACGAACAAGTGAAGGAGTGCAGCGACGGTCTGCTGTC<br>GACTATCGGCAAGAACCTGGACGATGACGCCGGTCGGATCGAAATCCGCGAGATGATGGCCAAATACTCGA<br>CCGACGTGATCGGCAGCTGCGCGTTTGGCCTGAAGCTGGATGCCATCAACGATCCGGACTCGGAGTTCCGG<br>AAGCACGGAAAAACCGTTTTTCAGCCGTCGCTGAGGTCCAAGATCCGAGTGGCGGTCATATTCATGCAGCC<br>GTCCCTGCTGAGCATTTTCCGCGTGCATCACTACTCGCACCGCACCATCCGATTCTTCCACGACGCGTTCC<br>AGCAGACGATTGAATATCGGGAAAAGCACAACGAAGACCGCAAAGACTTTGTGCAGCATCTGATGAAGGCC<br>AGAGAAGATCTAGTGTTGAACCCGAATCTAAAACCCGAAGAAAAATTCACTGAAATGGATATTGTAGCGAA<br>CGCATATATTCTCTTCATCGCTGGTTTCGAAACAGTATCTACATCAATGAGCTTTTGTATGTATGAATTAG<br>CATTAAGAAAAGATGTCCAAGATAAAGTTCGAAAAGAAATATTGGAAGTTAAATCTAAGTACAATGGACAA<br>ATGAATAGTGAATGCCTTAACGAACTTCATTATATGGGCATGGTTATTAAAGAAACATTGAGAAAATATCC<br>TCCATTAGTGACATTAAATCGAGTTGTGACTAAGCCGTATGTAATACCAGGGACACAAATCAAGTTAAAAA<br>TAGGTACTAAAATTGTTGTTCCAGTACATGCCATTCACTACGATCCAAAATATTACTCTGATCCAGAGGCT<br>TTTGAACCAGATCGTTTTTCAGATGAAAACATACATAATATACAACCTAACACATATATGCCTTTCGGAGA<br>CGGTCCTAGATTTTGTATTGGCAAACGATTTGCTGAATTCGAAATGAAAATGGCCTTGTCCGAAGTGTTAA<br>CCAACTACGAAGTGATGGCATGTGATAAAACCCAAATTCCCATAAAATATGTTATCGGAAGTTTTGTGAAT<br>ATACCTGAAAGCGTTTGGTTAAAATTTAGGAAAGTGAATACTTAA |
| ApRed | amplified from cDNA | ATGGAGAATCCTGAAGGAGAGAAAATTGAATCAACTGTTGTTTCAGAAGAGCCATTGATTAGTGCTTTAGA<br>TATTGGTCTTTTGGTGGTTATTATAACAGTTGGATATTTTTGGTACATCAAAAGAGATAAAAAGTCAAGTT<br>CTTCAGAAAAAAAACCCTATACTATTCAGCCATCTTCGTTGAGTTCTATTGAGCAGACCTCCAATAGTTCT<br>TTTATAAAAAAACTTAAATCAACCGGTCGTAGTTTAGTAGTGTTCTATGGAAGTCAAACAGGGACTGCAGA<br>AGAATTTGCTGGAAGAATAGCCAAAGAAGGGGCCAGATATAAAATGAAAGGAATGGTAGCCGATCCTGAAG<br>AATGTGATATGGCAGACTTGGTAGAAATGAAAGAAATACAAAAGTCATTGGCAATTTTTTGTATAGCAACA<br>TATGGTGAGGGAGACCCGACTGATAATGCTATGGATTTCTACGAATGGCTTCAAAATGGTGACGCCGACCT<br>AGAAGGATTGAATTATGCAGTATTTGGATTGGGAAATAAGACTTACGAGCATTATAACGAAATTGCTATTT<br>ACATTGATCAACGTTTGGAAGAATTGGGTGCTACTAGAGTTCATGAAATTGGTCTAGGAGATGATGATGCC<br>AATATTGAAGATGATTTTGTTTCTTGGAAAGAAAAATTATGGGATAGTGTTTGCTCCCATTATGGTATTGA<br>AGAAACTGGCGAAGAAAGTAATATTAGACAATACAAATTAGTAGACTGTTCCGAAGTTTTACCAGAGCGAA<br>TATTTTCGGGTGAAATATCACGACTTAAATCTTATGAAAACCAAAGATTCCCATTTGATGTTAAGAACCCA<br>TATTTATCTAAAATATCAGTTAATCGTGAGCTCCACAAGTCAGGTGATCGCTCTTGTATGCATATTGAGTT<br>TGATATTGATGGATCAAAGATGAGGTATGACACAGGAGATCATGTGGCTGTCTATCCCAAAAATTCTTCTG<br>AATTAGTAGAGAAAATTGGAGAATTGTTAAATGCAGATCTAGACACTGTATTTTCATTGTTGAACACTGAT<br>GAGGAATCCAGTAAAAAACATCCATTCCCTTGTCCTTGTACTTATCGAACAGCTTTAACTTATTATTTGGA<br>TATAACTTCAAATCCACGCACACATATTATGAAAGAATTAATAGAATATGCAAGTGATCCAAAGGATCAAG |

| | | |
|---|---|---|
| | | AAAAATTAAAGCTTATGGCAAGCTCAACCCCAGAGGGCAAAAAAGAATTTCACGAATGGATATTGCGTGAC<br>AACCGTAATATTGTTCATATTTTGGAAGATTTACCGAGTGTCAAACCAGATTTGGACCATTTATGTGAACT<br>ACTCCCACGATTGCAGTGTCGTTATTATTCAATATCCTCGTCACCAAAAGTGTACCCAAAATCTATTCATA<br>TTACAGCAGTTCTAGTAGAATACACTACTCCTACTAATCGAGTCAATAAAGGTGTTGCAACTAACTTATTA<br>GCCCAACTGAAACCAACCAATGACGAACTTCTACAACCTACTATACCTATTTATATTAGACGATCTCAATT<br>CAGATTGCCCCCTAAGAGTCAAACTCCAATTATAATGATTGGTCCCGGTACAGGATTAGCACCATTCCGAG<br>GTTTTATACAAGAAAGAGATTATGCACGTAAAGAAGGTAGAGAAATAGGAGAAATGGTTCTGTACTTTGGA<br>TGTCGAAAAAAGGATGAAGACTTCATTTATGAAAATGAGTTACAAGAATATGTTGCTAATGGAACCTTAAC<br>AAAGCTACATTTGGCATTTTCGCGTGATCAACCTGAAAAGCAGTATGTGACACATTTGTTAGAACAAAATG<br>CTGATGAACTGTGGAATATTATTGGTGAAAAAAATGGTCATTTATATGTTTGTGGTGATGCAAGGAGCATG<br>GCAAAGGATGTGCATAGTATTATTGAAAAAGTTGTCATGGAAAAGGGTCAAATGACCAATAGTCAGGCACT<br>CAATTATGTTAAAAAAATGGAACAACAGAAAAGATATTCTGCTGATGTGTGGAGCTGA |
| ApHGO | optimized for *E. coli* | ATGGAGAAATGGAATGGCAAAGTCGCTGTGGTCACAGGCGCCTCGAGTGGAATTGGCGAGGAGACCTGTCG<br>CCAGCTTGTGGAGCGCGGAATGATCGTTGTAGGGTTCGCACGCCGCGAGGACAAACTCCAGGAGCTGGAGA<br>AGGACTTAAAAGGGAAGCTGGGAAAGTTTTATTACGTGAAAGTAGATCTTTGCTCCGAGGAAAATATTATG<br>GAAGCATTCAACTGGGTAAAGAGCACTCTGAAATCGGTGGATGTTCTGGTCAATAATGCCGGCGTTCTGAA<br>GAAGAGTGATTTACTGGGCAACACTAACGATTGGAAACAGATGTTTGATACGAACGTTATGGGTCTGAATA<br>TTTGCTCACGTGAAGCCATTAAGATCATGGAAGAAATCCAAATTAAAGAGGGCCACATTATTAACATCAAT<br>AGTGTTGGTGGACACTACCAGTTCCAATTTGTAAAGGATTTCTCAGTCTACTGCGCCACGAAACACACCGT<br>GACCATCATTACAGAAAGCCTGCGCGAACTGATGGGTATGAAGAATCTTCCGGTCCGTGTTACGTCCATTA<br>GCCCGGGAGCGGTGGACACTGAGATGACTCTCGAGTTCTCAAAGATGGAAGGGTTCAAGATGCTTAAGAGC<br>ATCGACATTGCCGAGGCGATTTTATATGCCCTCAGCGCGCCGCAACGTGTAAACGTAGCGGAAATTATTAT<br>TCGTCCCACCGGCGAGAACACTGCGGGCTTGATTAAGAATTTCGTATGA |
| ApISY | amplified from cDNA | ATGATGGACGTCGTAAACGTTATGTTCGTCGGTATGACGTTCGTGTTCGTGGTTGTGCCGTTGGTGCTCAA<br>GTCGATGGACGGGACTCGATTGCCGGTCGCTGTGGCCCGATTGTACGCGTACGGCAAGATCACGGGCGGCG<br>CGAAGCCGTCCGGCGTGCTGAGCGTGCCCAAGCGGTGGTACAAGCACTTTTACGCATTCTCATTGGCCCTG<br>TCACTGATGGCAGCGCTGGCGCTGTCAGACGAGTATGCGGCGGGAGCCCCTTGGCCGTCGGCCGCGGCGTG<br>GTGCCGGCGGTGGCTGTGGGACCCGGTCCGCCGACCGAAGTCGTCGTACAGCTCGGCCGCCGCGTCCACGG<br>CCGCCGGCATGTTCGTGCTACAGTGCGCCAGGCGGACGTACGAGACGTTCCACGTGAACGTGTTCTCGGAC<br>ACGGCGGTTGGGCTGTGGTATTACGCGTCCGGTTACATGCACTACATAGGCGCCATCGTCACCGTGCTGGC<br>CGAGGCGCCGGTCGCAGTGGCGGCCGGGGAGCCACAGTCCAGCGGCGCCTGGGAGCCTGTCCGGTTGGCCG<br>GGGCCCTGCTGGTGTTTGCCTGGGCATATCGCGAACAGTGGCGGGCCAACGTGGCACTGGCCGAGGCCCGG<br>AAGCGCGGAGGTCAGGTGGTCACGCACGAGCACATTATGCTCACCGGCGGACTTTTCGACCTCGTGTCTAG<br>TCCACAGATGCTCACCGAGGTGGTCTTGTACGGCGCTTGGTACGCCGTGCTGTGGGGCACCACCGGATGGA<br>AGTACGTGATCGCGTTCGTCTGGGGAAATCAGTTCGAGATAGCACTCATCAGTCACCGGTGGTACCAGGAC<br>AAGTTCCCGAACTACCCACGAGAAAGGAAAGCTATCATACCGTATATTCTGTAA |
| ApNEPO | optimized for *E. coli* | ATGGCATCTCTTATCAGTGGGGCAATGAGCAGCGCAGCGTGGACCGGCGCGGGTATGATTCCTGTGTTCGT<br>GGTAGGACTTGCATACCTGCGTTATTCGATGTATGACCCGGAGTCCCGTGTGGTAGATGTGCTGGAAGTGC<br>GCGACGAGTACGACTTCATTGTGGTCGGCGCCGGTAGCGCAGGTGCAGTCATCGCAAATCGCTTGTCCGAA<br>ATGCAAAATTGGACCGTGTTAGTCTTAGAGGCCGGTGGTGACGAGACCGAAATTAGTGATGTTCCTTCTTT<br>CGTAGGCTATTTACAGCTTTCGGACATGGACTGGCAATATAAAACAGCCCCGCCGAGTAGTGATAATCCAT<br>ATTGCCTGGCGATGGTTCACGACCGTTGTAACTGGCCTCGTGGTAAGGTGCTGGGTGGCTCGTCGGTCTTG<br>AATGCAATGGTGTATGTCCGCGGTAATCAACGTGACTATGATATGTGGGCGGCCGCGGGAAATCCCGGTTG<br>GGCGTACGCTGACGTCTTGCCATATTTCCTCAAATCGGAAGATAATCGCAATCCGTACTTGGCGCGTACGA<br>AATATCATGCCCGTGGTGGCTACCTGACGGTCAGCGAGGCCCCGTGGCGCACGCCTCTCGCCACAGCTTTT<br>GTTGCGGCTGGTGAGGAGCTTGGTTACCAAAATCGCGATATCAACGGCCAATACCAAAATGGATTCATGCT<br>CACCCAAACGACGACCCGTCGTGGTTCCCGCTGTTCAACTGCCAAGGCGTTTCTGCGTCCTATTCGTCTGC<br>GTCCGAATATTCACGTGTCGATGCACAGTCAAGTAACTCGTATTCACTTCAGCGGTGGTAACGGCGGCAGC<br>GACAAACTTCGTGCAACAGGTGTCACTTACCTTCGCAATGGGAAACGCCGTACTGTTACTGCTCGCAAAGA<br>GGTAATCTTGAGTGCAGGGGCGATTGGCAGTCCCCAATTGCTGATGGTCAGCGGAGTAGGACCCGCTGATC<br>ACCTTACTGAACTGGGCATTAAGCCAGTCGTAGATCTGAAGGTAGGGCATAACTTACAGGATCACGTAGGC<br>TTAGGCGGTCTCACTTTCCTTATTGATGACCCAATCACGTTCAAGAAGTCACGCTTCACCTCAGCCTCGGT<br>GGCCCTGGATTACATCATGAACGAGCGTGGTCCTTTGACCTCGAGTGGAGTCGAGGGTCTCGCGTTCGTAA<br>ACACCAAGTACGCAGACCCGTCCGGCGAGTTTCCGGATATTCAGTTCCACTTTGCACCCTCGTCAGTCAAT<br>AGCGATGGGGACCAAATTCGCAAAATCACCGGTCTGCGCGACGCGGTTTACAATACAGTATACAAGCCATT<br>GGTGAATGCCGAAACATGGACTCTCCTTCCTCTCTTATTGCGTCCAAAGAGCTCGGGTTGGGTGCGCCTTA<br>AGAGTAAGAACCCACTTGCCCACCCCATCATCGAGCCCAATTACTTTGCCCACCGCGAGGACGTCCAAGTT<br>CTCGTCGACGGGATTCGCATCGCATTCAACGTAAGCAATACCGCCGCGTTTCGTAAGTACAATTCCCGTCC<br>ACTTTTAACCCCGATGCCAGGCTGTAAGAAGTTTGAGCTGTTCAGTGATGAGTACTGGGAATGCGCTCTGC<br>GCCACTTCACGTTTACCATCTACCACCCTGCGGGAACGTGTAAGATGGGCCCGGACACGGACCCCGACGCA<br>GTGGTTGACCACCGTTTACGCGTGCGTGGGATCGATGGCCTGCGTGTGATTGACGCCTCTATTATGCCTAA<br>TATCATCAGTGGTAATCCTAATGCCCCAGTGATCATGATTGGCGAGAAGGGTGCCGACATGATTAAGAAGG<br>ACTGGTCAAGCTAA |
| 100164798 | optimized for *E. coli* | ATGTCGGGAGTTGAAGTCATCCCAATTGGAGCAATCGCCGGAGCCGCCAGCCAGGTGGCCTGGTTTCTTCC<br>AGTTCTGGTTGCCGCCATCGCGTATTTCCATTATGAGGTCAACGATCCGGAGAGTCGCATTATTGATCAAC<br>CGGGCAATCTCATTTTAGACCAATACGATTTTATCATTGTGGGTGCAGGCAGCGCCGGTGCGGTCTTAGCA<br>AACCGTCTTACCGAGGTTGAGGACTGGTCGGTGCTTCTCATCGAAGCTGGCGGTGATGAGACGGAGATTTC<br>CGACGTGCCCCTTCTTGCCGCGTATTTGCAGTTGACTCAATTGGACTGGCAATATAAGGCGGAGCCCCAAG<br>ACACCGCCTGCTTAGCAATGAAAGATCAGCGCTGCAATTGGCCACGCGGTAAGGTTTTGGGCGGCAGTTCT<br>GTTTTGAACTATATGATGTACGTACGCGGTAATCAGATGGATTACGATAGTTGGTTAAAGCAAGGCAATCC<br>GGGCTGGGGTTACAACGACGTCCTTTATTATTTTAAGAAGTCTGAGGATAATCGTAACCCATATCTTGCTC |

| | | |
|---|---|---|
| | | GCACGCCCTACCACAGTATGGGCGGTTATTTAACAGTGTCCGAAGCACCGTATAAGACCCCTCTGGCAGAC<br>GCGTTTGTTCGTGCGGGCCAGGAGATGGGATACGACATCCGTGATATCAATGGCGAACGCCAGACAGGCTT<br>CATGATCCCTCAAGGCACCATCCGTCGCGGCGCCCGTTGCAGCACGGCAAAGGCCTTCTTACGCCCCGCCC<br>GTTTGCGTAAGAATTTACACGTCTCTATCAATGCCCACGTGACTCGCGTTGCGATTAACCCTGAGACCAAG<br>GTGGCATTTGGCGTGGAGATGATCAAGGACAACACGCGTCATTTCATTCGTGCAAATAAAGAGGTGTTACT<br>GTCCGCCGGCTCCATCTCATCGGCCCAACTTCTTATGCTGTCAGGCATTGGGCCGAAGAACCACCTTACTG<br>AAATGGGCATTCCGGTCTTGGCCGACTTAGATGTAGGAAAGAACCTGCAAGACCACGTAGGGTTAGGTGGC<br>TTGGCCTTTCTGATCAATAAGGAAGTTAGCTTGACCCAGGAACGCGTGGAAAATGTACAAACCGTGTTGAA<br>TTACGCCACCATGGGTGACGGACCGTTGACCGTCATGGGTGGCGTAGAGGGGTTAGCCTTTATCAATACAA<br>AATACGCGAACCAGAGCGTGGACACCCCTGATATTGAGCTCCACTTTGTTTCCGGAAGTACCAATAGTGAT<br>GGCGGTGTCCAATTATGGAAAGCCCACGGCTTAAAAGAGGAATTTTATAAGGCGGTGTACGAGCCAATCAA<br>TAACAAGGACGTGTGGTCTGCTATCCCCATGCTGTTGCGCCCGAAGAGCCGTGGAGAGATTCTTCTTCGCA<br>GCACCGACCCGTCCATGTACCCACGTATTCTGCCTAATTACCTCACCGTGCAAGAGGACGTCGATACCTTA<br>GTTGAGGGTGTGAAGTTTGTGGTGGCAATGTCTCGTACGACCCCATTCCGCCGTTATGGTTCCCGCCTGCA<br>CGATATTCCGTTCCCTGGTTGTGCGGCGGTGCCCCGTTTCACGGACGCGTATTGGGAGTGTATGGTCCGTC<br>ATTACACTGTGACCATCTATCATCCAGTGGGTACGGCAAAGATGGGCCCTGAGTGGGACAAGACAGCCGTA<br>GTGGACCCGCGCCTGCGCGTCTACGGTATTCACGGCCTTCGTGTAGTGGATGCCAGCATCATGCCCACCCT<br>TGTAAGCGCCAATACCAATGCACCTGTCATCATGATCGCTGAGAAGGCTGCTGACATGATCAAGGAGAAGT<br>GGCTGGGACGTAAGCGTTGA |
| 100168586 | optimized for *E. coli* | ATGAAGAAGATTGCGATTTTCGGGGCAACCGGTATGACAGGACTGTGTACCGTTGAGGCCGCATTAAAGCA<br>AGGGCTCGAGGTGCGCGCCTTGCTTCGTGACCCCAGCCGCATGCCCGAAGAGCTGCGCAAGCAGGTTGAGG<br>TAATCACAGGTGACGTCTTAGTGAAAGAGGATGTTGACAAGGTGGTTGAGGGCCGCGACGCGATCGTCGTC<br>ACACTGGGGACACGTAACGATTTAGCACCAACTACGATCATGTCAGAGGGTCTTCGTAATATTCTGTCGTC<br>AATGGAAAAGAACAATGTTAAGATCGTTAGCGTATGTCTTTCTACGTTCTTATTCTACGATAAACCCAAGG<br>TGCCCGCGATGTTCCATGGCATTAACGACGACCACGAGCGCATGTTACATCTCCTTCAGGCGGCAGAGAGC<br>CTGGACTGGATTGCTGTAATGCCGCCGCACATCGCGGGTACGCCGAGCGGGGACTACTCAGTTGAGATTGG<br>CTCAAGTCCGGGGCGTGCGATTAGCAAGTATGACTTGGGAAAGTTTATGATTGAGTGTCTTAGCAAACCAG<br>ATTATTATAAGCAACGCTGTGGTCTGGCTACGAAAGTTCCAGCGCCGTGA |
| 100162094 | optimized for *E. coli* | ATGGAGTACTTCCTGCCGAACCGCTGTACGTCAACTGTTCGTTTAGACGGCAAAACAGTCGTAGTTACTGG<br>CTGCAATACGGGCATTGGGAAGGAGACGGCAACGGAGTTCTACAAGCGCGGCGCACGCGTGATTATGGCTT<br>GTCGTTCGGCTAGCCGCACGCAAGATGCTATTGAATCGATCAAGAATCAAACGGAAGGCGATAACAACGTG<br>GGTGAGTTGGTCTTTAAGCACTTGGAGCTTAGCTTTCTGGCATCCGTACGTAAGTGCGCGAAGGAGATCCT<br>GCACACGGAGAAACGCATCGACATTTTAGTCAACAACGCCGGCATTATGATGTGCCCCAAGACGCTTAGTG<br>AGAATGGTATCGAGCTGCATTTGGCCACTAATCATCTGGGCCACTTCCTTTTCACTCTGTTGTTATTACCG<br>CGCATCCTGAAGAGCGCGCCCGCACGCATTATCAACGTCACATCCCTGGCTCACAAATGGGGAGATCAGAA<br>GATGCACTTTGATGACATTAATCTTGATAAGGACTACACCCCGTCAGGCGCGTACGGACGCTCAAAGCTCG<br>CCAATATCCTCTTTACGGTGGAGCTTGCTAAACGCCTGAACGGGACGGGCGTAACGGTTTACGCTGTAAAT<br>CCCGGTATTGTACATACCGAGCTCTCGCGTTATGTCGACCAAACTATTTTCCCAGGTGCGTCCTGGCTTTA<br>TAACTCATTCACGAAGATTGCAGTGAAGACTCCTCAACAAGGCGCACAAACGACCTTGCACTGTGCGCTTG<br>ATGAGAAATGTGCTGGCGAAAGCGGCTTGTACTACAGTGACTGTAAGGTTCTTGAGCCTGAGCCAGTAGCA<br>AAAGACGAAGAAGTCAGCGCCCAATTGTGGGATACTTCCTGCGCTTTTGTTAACCTGGAACCGTCTATTGA<br>TCCTTTTAAGCCCGAATCGGACGATGTCTCCATTTGA |
| 100162683 | optimized for *E. coli* | ATGTTAAGCCGTCGCTGCGCATATCGCTCGCAGGTTTCTCGTGGCGCGCACGGCGTTTGCACGATGTTTAG<br>CTCGGATGGCATGACATGTCCTCCTGATGCTGATGGTACGGCAGCCTGTGGCCATCGGCGTCGCTGCTCAG<br>GTCTGATCGGTTGCGCATTGCTCGCCTCTGAGGCCGCTGCGCACGTCGCTCGTCTTTGTCGCTCAAATCCG<br>CGCCTGCTGTCCATGCTGGTGCAGGAGAAATGCGGTGATGATGCCAATAGTTCACGCTTTGCTCATGATTT<br>CAAAACCCTGGCTGATGTTCTGGTTCAACGCGTGGTATCTAAACGTATTGGCCGCCAATTTCCGGAGTTAG<br>AAGGTAACGTGTATGGGGAGGAAAATGATACCATTCAGAATAATAAAGGCCAAGACGTCACTATTCAGGTT<br>GGCGAAACGGTTGAGGAAACGCTGCAGTGCCTTACTAAAGCGCTCGGCGATGACTTGGACAGTGCCAAGAG<br>CTTGGCCGAAGCGGCCCATAAAGAGTTTGAAAGTAAGGATCTCAACGTGGATTGCTATCCGCCAGAAGAAG<br>AGAACTTAGATCTGGAAAAAATCGGGATTTGGATCGACCCGATTGATTCGACCAACGAGTATATCAATGGA<br>AATGTGGACACCATCAATGAATATGGCTTTCATTCGTCAGGCCTGCATTGCGTGACTGTGAACATTGGTCT<br>GTTCGATAAATATTCTGGTAAACCCATTGCGGGGGTGATTAATCAGCCTTTCTTCGAATTCGACGCGGTCA<br>ACGGCTGGAACGGCCGCTGTTACTGGGCATACTGCGACGGCCAGAAATCGCTGAACAGTCTCCCAGAATTT<br>ATTAGCTGTAATCAGGAATTAGTGGTCACTTCCAACAGCGAAACTGATGCGGCTAAAGCCGCATTACGTCG<br>TTCCGGTTATACCGTTGCAACCGCGAGTGGAGCGGGCTATAAAATGCTTTGTGTGGCGCTGGGAATCGTGA<br>AATGTTATGCGCTTACCAAAGACTCAACGTACGCCTGGGACACCTGCGCGGCTCATGCTATGTTGGCGTCG<br>CAGGGCGGAAAAGCCTGTCAGTGTAACACAGCAGATGGGCCTTTAACTTATCGTCCGAAAACTGCGGGTGG<br>CGGTCGGGCCCATTGCAATGCAGCCGGCGTGATTGCTTCCCGTGATCCACAAACGGTTGACCGTGTCCATG<br>CCCTGTTATCCCCGTTACATCTGCATTGCAGCAGTAGTTCATAA |

**Table S5: Prediction of transmembrane domains by DeepTMHMM.** S, signal peptide; I, inside cell/cytosol; M, alpha membrane; B, beta membrane; P, periplasm; O, outside cell/lumen of ER/Golgi/lysosomes.

| Protein | Transmembrane domain prediction |
| --- | --- |
| ApIDS | MNKMLTFTRALSRRSAFLLSDSAAVRENCFRSMSTVRAPPVPPVITGTAVSKDETRDFMAVFPDVVRDLTDTGRNLDVPD<br>VTKWLAKLLQYNVPGGKKNRGLALVLSYKMLSSPADQTDENIRLSYILGWCVEILQAYQLVLDDIMDNAITRRGRPCWYR<br>HNDIGLMAVNDGVLLEQSIYQLIKKYFKDKPYYTHILELFYDVTMKTSMGQCLDMLTANSFKSKKLEKYTMENYTAIVKY<br>KTAYYSFFLPVCLAMRMTNINDPEIFRQAKTILLEMGHFFQVQDDFLDCYGDPDVMGKIGTDIEDGKCSWLAVVALQKVN<br>SEQKKIMEDNYGIDNPANVAVIKDLYAQLKLPDTFHLYEEESYKLICTHIQQLSRGLSQDMFFKFLEKIYKRTL<br>IIIIIIIIIIIIIIIIIIIIIIIIIIIIIIIIIIIIIIIIIIIIIIIIIIIIIIIIIIIIIIIIIIIIIIIIIIIIIIII<br>IIIIIIIIIIIIIIIIIIIIIIIIIIIIIIIIIIIIIIIIIIIIIIIIIIIIIIIIIIIIIIIIIIIIIIIIIIIIIIII<br>IIIIIIIIIIIIIIIIIIIIIIIIIIIIIIIIIIIIIIIIIIIIIIIIIIIIIIIIIIIIIIIIIIIIIIIIIIIIIIII<br>IIIIIIIIIIIIIIIIIIIIIIIIIIIIIIIIIIIIIIIIIIIIIIIIIIIIIIIIIIIIIIIIIIIIIIIIIIIIIIII<br>IIIIIIIIIIIIIIIIIIIIIIIIIIIIIIIIIIIIIIIIIIIIIIIIIIIIIIIIIIIIIIIIIIIIIIIIII<br># ApIDS Number of predicted TMRs: 0 |
| ApGES | MMTYSVSSNKKHDLSNRNIFGKLLALFSLTPFVILSGFISLILFRRDLHTITFFFGVLLNEICNTVLKHILREPRPLARN<br>TNLLYSEYGMPSSHSQFMWFFASYMLYFTFIRLQYANNKAFKEFFWKVAGAVSCIAIACIVSYSRIFLQYHTWKQVIYGA<br>LFGIIIGTIWFTIINVVLTPYFPTVISWKISELFLLRDTTLIPNVLWFEYTNIRHEAGARARRRKSISAKSQ<br>OOOOOOOOOOOOOOOOOOOOOOOMMMMMMMMMMMMMMMMMMMMMIIIIMMMMMMMMMMMMMMMMMMMOOOOOOOOOOOOO<br>OOOOOOOOOOOOOOMMMMMMMMMMMMMMMMMIIIIIIIIIIIIIMMMMMMMMMMMMMMMMMMMMMMOOOOOOOOMMMMMM<br>MMMMMMMMMMMMMMMIIIIIIIIIIIIIIIIIIIIIIIIIIIIIIIIIIIIIIIIIIIIIIIIIIIIIIIIII<br># ApGES Number of predicted TMRs: 5 |
| ApG8H | MFEFVYELFDLKMLLVTAFLGAIYVYSTWTHSHWSKLGISSPSAPVPLFGHAMPSMLGQMHFMDVLHNLYKELGDQRFGG<br>IYTMRTPQLLVKDPELIGHILIKDFNNFTDRGLYAGTHTNPLNNNIFFTRGERWKTMRQKLSPTFTANKLKYMNEQVKEC<br>SDGLLSTIGKNLDDDAGRIEIREMMAKYSTDVIGSCAFGLKLDAINDPDSEFRKHGKTVFQPSLRSKIRVAVIFMQPSLL<br>SIFRVHHYSHRTIRFFHDAFQQTIEYREKHNEDRKDFVQHLMKAREDLVLNPNLKPEEKFTEMDIVANAYILFIAGFETV<br>STSMSFCMYELALRKDVQDKVRKEILEVKSKYNGQMNSECLNELHYMGMVIKETLRKYPPLVTLNRVVTKPYVIPGTQIK<br>LKIGTKIVVPVHAIHYDPKYYSDPEAFEPDRFSDENIHNIQPNTYMPFGDGPRFCIGKRFAEFEMKMALSEVLTNYEVMA<br>CDKTQIPIKYVIGSFVNIPESVWLKFRKVNT<br>IIIIIIIIIIIIIIIIIIIIIIIIIIIIIIIIIIIIIIIIIIIIIIIIIIIIIIIIIIIIIIIIIIIIIIIIIIIIIIII<br>IIIIIIIIIIIIIIIIIIIIIIIIIIIIIIIIIIIIIIIIIIIIIIIIIIIIIIIIIIIIIIIIIIIIIIIIIIIIIIII<br>IIIIIIIIIIIIIIIIIIIIIIIIIIIIIIIIIIIIIIIIIIIIIIIIIIIIIIIIIIIIIIIIIIIIIIIIIIIIIIII<br>IIIIIIIIIIIIIIIIIIIIIIIIIIIIIIIIIIIIIIIIIIIIIIIIIIIIIIIIIIIIIIIIIIIIIIIIIIIIIIII<br>IIIIIIIIIIIIIIIIIIIIIIIIIIIIIIIIIIIIIIIIIIIIIIIIIIIIIIIIIIIIIIIIIIIIIIIIIIIIIIII<br>IIIIIIIIIIIIIIIIIIIIIIIIIIIIIIIIIIIIIIIIIIIIIIIIIIIIIIIIIIIIIIIIIIIIIIIIIIIIIIII<br>IIIIIIIIIIIIIIIIIIIIIIIIIIIIIII<br># ApG8H Number of predicted TMRs: 0 |
| ApRed | MENPEGEKIESTVVSEEPLISALDIGLLVVIITVGYFWYIKRDKKSSSSEKKPYTIQPSSLSSIEQTSNSSFIKKLKSTG<br>RSLVVFYGSQTGTAEEFAGRIAKEGARYKMKGMVADPEECDMADLVEMKEIQKSLAIFCIATYGEGDPTDNAMDFYEWLQ<br>NGDADLEGLNYAVFGLGNKTYEHYNEIAIYIDQRLEELGATRVHEIGLGDDDANIEDDFVSWKEKLWDSVCSHYGIEETG<br>EESNIRQYKLVDCSEVLPERIFSGEISRLKSYENQRFPFDVKNPYLSKISVNRELHKSGDRSCMHIEFDIDGSKMRYDTG<br>DHVAVYPKNSSELVEKIGELLNADLDTVFSLLNTDEESSKKHPFPCPCTYRTALTYYLDITSNPRTHIMKELIEYASDPK<br>DQEKLKLMASSTPEGKKEFHEWILRDNRNIVHILEDLPSVKPDLDHLCELLPRLQCRYYSISSSPKVYPKSIHITAVLVE<br>YTTPTNRVNKGVATNLLAQLKPTNDELLQPTIPIYIRRSQFRLPPKSQTPIIMIGPGTGLAPFRGFIQERDYARKEGREI<br>GEMVLYFGCRKKDEDFIYENELQEYVANGTLTKLHLAFSRDQPEKQYVTHLLEQNADELWNIIGEKNGHLYVCGDARSMA<br>KDVHSIIEKVVMEKGQMTNSQALNYVKKMEQQKRYSADVWS<br>OOOOOOOOOOOOOOOOOOOOOMMMMMMMMMMMMMMMMMIIIIIIIIIIIIIIIIIIIIIIIIIIIIIIIIIIIIIIIIII<br>IIIIIIIIIIIIIIIIIIIIIIIIIIIIIIIIIIIIIIIIIIIIIIIIIIIIIIIIIIIIIIIIIIIIIIIIIIIIIIII<br>IIIIIIIIIIIIIIIIIIIIIIIIIIIIIIIIIIIIIIIIIIIIIIIIIIIIIIIIIIIIIIIIIIIIIIIIIIIIIIII<br>IIIIIIIIIIIIIIIIIIIIIIIIIIIIIIIIIIIIIIIIIIIIIIIIIIIIIIIIIIIIIIIIIIIIIIIIIIIIIIII<br>IIIIIIIIIIIIIIIIIIIIIIIIIIIIIIIIIIIIIIIIIIIIIIIIIIIIIIIIIIIIIIIIIIIIIIIIIIIIIIII<br>IIIIIIIIIIIIIIIIIIIIIIIIIIIIIIIIIIIIIIIIIIIIIIIIIIIIIIIIIIIIIIIIIIIIIIIIIIIIIIII<br>IIIIIIIIIIIIIIIIIIIIIIIIIIIIIIIIIIIIIIIIIIIIIIIIIIIIIIIIIIIIIIIIIIIIIIIIIIIIIIII<br>IIIIIIIIIIIIIIIIIIIIIIIIIIIIIIIIIIIIIIIIIIIIIIIIIIIIIIIIIIIIIIIIIIIIIIIIIIIIIIII<br>IIIIIIIIIIIIIIIIIIIIIIIIIIIIIIIIIIIIIIIII<br># ApRed Number of predicted TMRs: 1 |
| ApHGO | MEKWNGKVAVVTGASSGIGEETCRQLVERGMIVVGFARREDKLQELEKDLKGKLGKFYYVKVDLCSEENIMEAFNWVKST<br>LKSVDVLVNNAGVLKKSDLLGNTNDWKQMFDTNVMGLNICSREAIKIMEEIQIKEGHIININSVGGHYQFQFVKDFSVYC<br>ATKHTVTIITESLRELMGMKNLPVRVTSISPGAVDTEMTLEFSKMEGFKMLKSIDIAEAILYALSAPQRVNVAEIIIRPT<br>GENTAGLIKNFV<br>IIIIIIIIIIIIIIIIIIIIIIIIIIIIIIIIIIIIIIIIIIIIIIIIIIIIIIIIIIIIIIIIIIIIIIIIIIIIIIII<br>IIIIIIIIIIIIIIIIIIIIIIIIIIIIIIIIIIIIIIIIIIIIIIIIIIIIIIIIIIIIIIIIIIIIIIIIIIIIIIII<br>IIIIIIIIIIIIIIIIIIIIIIIIIIIIIIIIIIIIIIIIIIIIIIIIIIIIIIIIIIIIIIIIIIIIIIIIIIIIIIII<br>IIIIIIIIIIII<br># ApHGO Number of predicted TMRs: 0 |
| ApISY | MMDVVNVMFVGMTFVFVVVPLVLKSMDGTRLPVAVARLYAYGKITGGAKPSGVLSVPKRWYKHFYAFSLALSLMAALALS<br>DEYAAGAPWPSAAAWCRRWLWDPVRRPKSSYSSAAASTAAGMFVLQCARRTYETCHVNVFSDTAVGLWYYASGYMHYIGA<br>IVTVLAEAPVAVAAGEPQSSGAWEPVRLAGALLVFAWAYREQWRANVALAEARKRGGQVVTHEHIMLTGGLFDLVSSPQM<br>LTEVVLYGAWYAVLWGTTGWKYVIAFVWGNQFEIALISHQWYQDKFPNYPRERKAIIPYIL |

| | |
|---|---|
| | OOOOMMMMMMMMMMMMMMMMMMMIIIIIIIIIIIIIIIIIIIIIIIIIIIIIIIIIIIMMMMMMMMMMMMMMMMMMMMMM<br>OOOOOOOOOOOOOOOOOOOOOOOOOOOOOOOOOMMMMMMMMMMMMMMMMMMMIIIIIIIIIIIIIMMMMMMMMMMMMMMM<br>MMMMMMMOOOOOOOOOOOOOOOOOOOOMMMMMMMMMMMMIIIIIIIIIIIIIIIIIIIIIIIIIIIIIIIIIIIIIIIIM<br>MMMMMMMMMMMMMMOOOMMMMMMMMMMMMMMMMMMMMMIIIIIIIIIIIIIIIIIIIIIII<br># ApISY Number of predicted TMRs: 7 |
| ApNEPO | MASLISGAMSSAAWTGAGMIPVFVVGLAYLRYSMYDPESRVVDVLEVRDEYDFIVVGAGSAGAVIANRLSEMQNWTVLVL<br>EAGGDETEISDVPSFVGYLQLSDMDWQYKTAPPSSDNPYCLAMVHDRCNWPRGKVLGGSSVLNAMVYVRGNQRDYDMWAA<br>AGNPGWAYADVLPYFLKSEDNRNPYLARTKYHARGGYLTVSEAPWRTPLATAFVAAGEELGYQNRDINGQYQNGFMLTQT<br>TTRRGSRCSTAKAFLRPIRLRPNIHVSMHSQVTRIHFSGGNGGSDKLRATGVTYLRNGKRRTVTARKEVILSAGAIGSPQ<br>LLMVSGVGPADHLTELGIKPVVDLKVGHNLQDHVGLGGLTFLIDDPITFKKSRFTSASVALDYIMNERGPLTSSGVEGLA<br>FVNTKYADPSGEFPDIQFHFAPSSVNSDGDQIRKITGLRDAVYNTVYKPLVNAETWTLLPLLLRPKSSGWVRLKSKNPLA<br>HPIIEPNYFAHREDVQVLVDGIRIAFNVSNTAAFRKYNSRPLLTPMPGCKKFELFSDEYWECALRHFTFTIYHPAGTCKM<br>GPDTDPDAVVDHRLRVRGIDGLRVIDASIMPNIISGNPNAPVIMIGEKGADMIKKDWSS<br>SSSSSSSSSSSSSSSSSSSSSSSSSSSSSSSSOOOOOOOOOOOOOOOOOOOOOOOOOOOOOOOOOOOOOOOOOOOOOOOO<br>OOOOOOOOOOOOOOOOOOOOOOOOOOOOOOOOOOOOOOOOOOOOOOOOOOOOOOOOOOOOOOOOOOOOOOOOOOOOOOOO<br>OOOOOOOOOOOOOOOOOOOOOOOOOOOOOOOOOOOOOOOOOOOOOOOOOOOOOOOOOOOOOOOOOOOOOOOOOOOOOOOO<br>OOOOOOOOOOOOOOOOOOOOOOOOOOOOOOOOOOOOOOOOOOOOOOOOOOOOOOOOOOOOOOOOOOOOOOOOOOOOOOOO<br>OOOOOOOOOOOOOOOOOOOOOOOOOOOOOOOOOOOOOOOOOOOOOOOOOOOOOOOOOOOOOOOOOOOOOOOOOOOOOOOO<br>OOOOOOOOOOOOOOOOOOOOOOOOOOOOOOOOOOOOOOOOOOOOOOOOOOOOOOOOOOOOOOOOOOOOOOOOOOOOOOOO<br>OOOOOOOOOOOOOOOOOOOOOOOOOOOOOOOOOOOOOOOOOOOOOOOOOOOOOOOOOOOOOOOOOOOOOOOOOOOOOOOO<br>OOOOOOOOOOOOOOOOOOOOOOOOOOOOOOOOOOOOOOOOOOOOOOOOOOOOOOOOO<br># ApNEPO Number of predicted TMRs: 0 |
| 100164798 | MSGVEVIPIGAIAGAASQVAWFLPVLVAAIAYFHYEVNDPESRIIDQPGNLILDQYDFIIVGAGSAGAVLANRLTEVEDW<br>SVLLIEAGGDETEISDVPLLAAYLQLTQLDWQYKAEPQDTACLAMKDQRCNWPRGKVLGGSSVLNYMMYVRGNQMDYDSW<br>LKQGNPGWGYNDVLYYFKKSEDNRNPYLARTPYHSMGGYLTVSEAPYKTPLADAFVRAGQEMGYDIRDINGERQTGFMIP<br>QGTIRRGARCSTAKAFLRPARLRKNLHVSINAHVTRVAINPETKVAFGVEMIKDNTRHFIRANKEVLLSAGSISSAQLLM<br>LSGIGPKNHLTEMGIPVLADLDVGKNLQDHVGLGGLAFLINKEVSLTQERVENVQTVLNYATMGDGPLTVMGGVEGLAFI<br>NTKYANQSVDTPDIELHFVSGSTNSDGGVQLWKAHGLKEEFYKAVYEPINNKDVWSAIPMLLRPKSRGEILLRSTDPSMY<br>PRILPNYLTVQEDVDTLVEGVKFVVAMSRTTPFRRYGSRLHDIPFPGCAAVPRFTDAYWECMVRHYTVTIYHPVGTAKMG<br>PEWDKTAVVDPRLRVYGIHGLRVVDASIMPTLVSANTNAPVIMIAEKAADMIKEKWLGRKR<br>SSSSSSSSSSSSSSSSSSSSSSSSSSSSSSSSOOOOOOOOOOOOOOOOOOOOOOOOOOOOOOOOOOOOOOOOOOOOOOOO<br>OOOOOOOOOOOOOOOOOOOOOOOOOOOOOOOOOOOOOOOOOOOOOOOOOOOOOOOOOOOOOOOOOOOOOOOOOOOOOOOO<br>OOOOOOOOOOOOOOOOOOOOOOOOOOOOOOOOOOOOOOOOOOOOOOOOOOOOOOOOOOOOOOOOOOOOOOOOOOOOOOOO<br>OOOOOOOOOOOOOOOOOOOOOOOOOOOOOOOOOOOOOOOOOOOOOOOOOOOOOOOOOOOOOOOOOOOOOOOOOOOOOOOO<br>OOOOOOOOOOOOOOOOOOOOOOOOOOOOOOOOOOOOOOOOOOOOOOOOOOOOOOOOOOOOOOOOOOOOOOOOOOOOOOOO<br>OOOOOOOOOOOOOOOOOOOOOOOOOOOOOOOOOOOOOOOOOOOOOOOOOOOOOOOOOOOOOOOOOOOOOOOOOOOOOOOO<br>OOOOOOOOOOOOOOOOOOOOOOOOOOOOOOOOOOOOOOOOOOOOOOOOOOOOOOOOOOOOOOOOOOOOOOOOOOOOOOOO<br>OOOOOOOOOOOOOOOOOOOOOOOOOOOOOOOOOOOOOOOOOOOOOOOOOOOOOOOOOOOOO<br># ApGMC-Ox2 Number of predicted TMRs: 0 |
| 100168586 | MKKIAIFGATGMTGLCTVEAALKQGLEVRALLRDPSRMPEELRKQVEVITGDVLVKEDVDKVVEGRDAIVVTLGTRNDLA<br>PTTIMSEGLRNILSSMEKNNVKIVSVCLSTFLFYDKPKVPAMFHGINDDHERMLHLLQAAESLDWIAVMPPHIAGTPSGD<br>YSVEIGSSPGRAISKYDLGKFMIECLSKPDYYKQRCGLATKVPAP<br>IIIIIIIIIIIIIIIIIIIIIIIIIIIIIIIIIIIIIIIIIIIIIIIIIIIIIIIIIIIIIIIIIIIIIIIIIIIIIIII<br>IIIIIIIIIIIIIIIIIIIIIIIIIIIIIIIIIIIIIIIIIIIIIIIIIIIIIIIIIIIIIIIIIIIIIIIIIIIIIIII<br>IIIIIIIIIIIIIIIIIIIIIIIIIIIIIIIIIIIIIIIIIIIII<br># 100168586_Flavin_reductase Number of predicted TMRs: 0 |
| 100162094 | MEYFLPNRCTSTVRLDGKTVVVTGCNTGIGKETATEFYKRGARVIMACRSASRTQDAIESIKNQTEGDNNVGELVFKHLE<br>LSFLASVRKCAKEILHTEKRIDILVNNAGIMMCPKTLSENGIELHLATNHLGHFLFTLLLLPRILKSAPARIINVTSLAH<br>KWGDQKMHFDDINLDKDYTPSGAYGRSKLANILFTVELAKRLNGTGVTVYAVNPGIVHTELSRYVDQTIFPGASWLYNSF<br>TKIAVKTPQQGAQTTLHCALDEKCAGESGLYYSDCKVLEPEPVAKDEEVSAQLWDTSCAFVNLEPSIDPFKPESDDVSI<br>IIIIIIIIIIIIIIIIIIIIIIIIIIIIIIIIIIIIIIIIIIIIIIIIIIIIIIIIIIIIIIIIIIIIIIIIIIIIIIII<br>IIIIIIIIIIIIIIIIIIIIIIIIIIIIIIIIIIIIIIIIIIIIIIIIIIIIIIIIIIIIIIIIIIIIIIIIIIIIIIII<br>IIIIIIIIIIIIIIIIIIIIIIIIIIIIIIIIIIIIIIIIIIIIIIIIIIIIIIIIIIIIIIIIIIIIIIIIIIIIIIII<br>IIIIIIIIIIIIIIIIIIIIIIIIIIIIIIIIIIIIIIIIIIIIIIIIIIIIIIIIIIIIIIIIIIIIIIIIIIIIIII<br># 100162094_retinol_dehydrogenase Number of predicted TMRs: 0 |
| 100162683 | MLSRRCAYRSQVSRGAHGVCTMFSSDGMTCPPDADGTAACGHRRRCSGLIGCALLASEAAAHVARLCRSNPRLLSMLVQE<br>KCGDDANSSRFAHDFKTLADVLVQRVVSKRIGRQFPELEGNVYGEENDTIQNNKGQDVTIQVGETVEETLQCLTKALGDD<br>LDSAKSLAEAAHKEFESKDLNVDCYPPEEENLDLEKIGIWIDPIDSTNEYINGNVDTINEYGFHSSGLHCVTVNIGLFDK<br>YSGKPIAGVINQPFFEFDAVNGWNGRCYWAYCDGQKSLNSLPEFISCNQELVVTSNSETDAAKAALRRSGYTVATASGAG<br>YKMLCVALGIVKCYALTKDSTYAWDTCAAHAMLASQGGKACQCNTADGPLTYRPKTAGGGRAHCNAAGVIASRDPQTVDR<br>VHALLSPLHLHCSSSS<br>IIIIIIIIIIIIIIIIIIIIIIIIIIIIIIIIIIIIIIIIIIIIIIIIIIIIIIIIIIIIIIIIIIIIIIIIIIIIIIII<br>IIIIIIIIIIIIIIIIIIIIIIIIIIIIIIIIIIIIIIIIIIIIIIIIIIIIIIIIIIIIIIIIIIIIIIIIIIIIIIII<br>IIIIIIIIIIIIIIIIIIIIIIIIIIIIIIIIIIIIIIIIIIIIIIIIIIIIIIIIIIIIIIIIIIIIIIIIIIIIIIII<br>IIIIIIIIIIIIIIIIIIIIIIIIIIIIIIIIIIIIIIIIIIIIIIIIIIIIIIIIIIIIIIIIIIIIIIIIIIIIIIII<br>IIIIIIIIIIIIIIIIIIIIIIIIIIIIIIIIIIIIIIIIIIIIIIIIIIIIIIIIIIIIIIIIIIIIIIIIIIIIIIII<br>IIIIIIIIIIIIIIII<br># 100162683_inositol_polyphosphate_1-phosphatase Number of predicted TMRs: 0 |

**Table S6: Primers used in this study.**

| Gene | Direction | Note | Sequence |
|---|---|---|---|
| *ApRed* | fwd | *Bam*HI | AGAGGGATCCGTAATGGAGAATCCTGAAGGAGA |
| *ApRed* | rev | *Xho*I | CGAGCTCGAGTCAGCTCCACACATCAGC |
| *ApG8H* | fwd | *Not*I | CCAGGCGGCCGCAATGTTTGAATTCGTCTACGAAC |
| *ApG8H* | rev | *Sac*I | CCAAGAGCTCTTAAGTATTCACTTTCCTAAATTTTAACC |
| *ApGES* | fwd | *Not*I | CGTGCGGCCGCAATGATGACTTACTCCGTTTCT |
| *ApGES* | rev | *Sac*I | ACAGAGCTCTTATTGTGACTTGGCGGAA |
| *ApISY* | fwd | *Not*I | GCAGCGGCCGCAATGATGGACGTCGTAAACG |
| *ApISY* | rev | *Sac*I | TCGGAGCTCTTACAGAATATACGGTATGATAGCT |

**Table S7: The iridoid pathway genes are randomly distributed throughout the *A. pisum* genome.**

| Gene | Scaffold (JIC 1.1.0) | Position |
|---|---|---|
| *ApISY* | 2 | 63,420,766 |
| *ApHGO* | 2 | 11,700,000 |
| *ApG8H* | 2 | 98,800,000 |
| *ApIDS* | 3 | 81,800,000 |
| *ApGES* | 1 | 127,500,000 |
| *ApNEPO* | 4 | 36,000,000 |
